# Distinct maternofetal immune signatures delineate preterm birth onset following urinary tract infection

**DOI:** 10.1101/2024.10.22.619711

**Authors:** Samantha Ottinger, Addison B. Larson, Vicki Mercado-Evans, Holly Branthoover, Jacob J. Zulk, Camille Serchejian, Marlyd E. Mejia, Zainab A. Hameed, Ryan Walde, Rachel C. Fleck, Allyson E. Shea, Kathryn A. Patras

## Abstract

Preterm birth is the leading cause of infant mortality resulting in over one million neonatal deaths annually. Maternal urinary tract infection (UTI) during pregnancy increases risk for preterm birth; however, biological processes mediating UTI-associated preterm birth are not well-described. We established a murine maternal UTI model in which challenge with uropathogenic *E. coli* resulted in preterm birth in about half of dams. Dams experiencing preterm birth displayed excessive bladder inflammation and altered uteroplacental T cell polarization compared to non-laboring infected dams, with no differences in bacterial burdens. Additional factors associated with preterm birth included higher proportions of male fetuses and lower maternal serum IL-10. Furthermore, exogenous maternal IL-10 treatment absolved UTI-associated preterm birth but contributed to fetal growth restriction in this model. Using urine samples from a cohort of human pregnancies with or without UTI, we correlated urinary cytokines with birth outcomes and urine culture status. These analyses yielded a non-invasive, highly predictive three-model system for evaluating preterm birth risk implicating cytokines IL-10, IL-15, IL-1β, and IL-1RA. Our unique bimodal murine model coupled with patient samples provides a platform to investigate immunological and microbial factors governing UTI-associated preterm birth, revealing novel therapeutic opportunities to predict or prevent preterm birth.

## INTRODUCTION

Preterm birth is the leading cause of neonatal mortality, impacting 15 million births and resulting in 1 million deaths annually(1). Surviving infants are subject to physical, neurodevelopmental, and socioeconomic sequelae. Up to 40% of preterm births are attributable to maternal or fetal infection, including urinary tract infection (UTI)(2–4). Of note, maternal UTI is highly prevalent, affecting 1 in 10 pregnancies(5). Several maternal factors, including physiological, hormonal, and socioeconomic aspects, may contribute to higher UTI incidence(6). Maternal immunological adaptation to support the fetus during pregnancy may also elevate UTI risk(7–9); however, the impact of pregnancy on bladder immunity has not been described.

UTI in pregnancy is associated with increased risk of adverse outcomes including preterm birth, intrauterine growth restriction, stillbirth, and neonatal sepsis(4, 10, 11). Despite abundant clinical correlations, there is limited data exploring this relationship mechanistically, in part due to lack of animal models that mimic clinical presentation in humans. A mouse model of maternal pyelonephritis with uropathogenic *Escherichia coli* (UPEC), the cause of 70% of UTIs, resulted in preterm birth in 90% of dams; however, this model found extensive placental and fetal bacterial invasion and maternal bacteremia and may not represent the impact of localized bladder inflammation(12). An outbred cystitis model observed increased uteroplacental immune infiltration upon infection and intrauterine growth restriction in pups born to infected dams; however, no preterm birth was detected(13). Furthermore, UTI has recently been shown to induce neutrophilic inflammation in mammary tissue of lactating mice(14). These latter studies support the role of localized bladder infection on distal tissue inflammation at sites critical for maternal-neonatal health.

Immunological tolerance during pregnancy, essential to prevent rejection of the semi-allogeneic fetus, is driven by type 2 and regulatory T helper cells(9). At parturition, inflammatory immune cells, such as cytotoxic and type 17 T helper cells, neutrophils, and macrophages, infiltrate the uteroplacental space, where they secrete interleukin (IL)-1β, IL-6, and matrix metalloproteinases that contribute to uterine contractility and fetal membrane rupture(15). Untimely recruitment of these cells has been implicated in infection-associated and spontaneous preterm birth(15). Sex-specific placental immune responses have been described in animal models and clinical studies, with heightened inflammation in males offspring(16–18). This is consistent with worse clinical outcomes for male neonates, with notable increased risk of preterm birth and neonatal morbidity(1, 19, 20).

Here, we describe a mouse model of UTI-associated preterm birth. This model recapitulates clinical manifestations of maternal UTI including preterm birth and intrauterine growth restriction in a subset of individuals, allowing us to investigate the impact of localized maternal inflammation on reproductive physiology and outcomes. Using this model, we describe immunological differences in the pregnant and non-pregnant bladder. We further find disparate bladder, systemic, and uteroplacental inflammation in preterm and non-preterm infected dams during acute UTI. These findings are supported by a cross-sectional human cohort of pregnant and non-pregnant UTI, in which we identify urinary biomarkers associated with preterm birth and low birth weight. These findings advance our understanding of how pregnancy modulates the immune response to UTI, while identifying biomarkers and therapeutic opportunities to predict or prevent UTI-associated preterm birth.

## RESULTS

### Maternal UTI induces preterm labor and intrauterine growth restriction

To establish a model of maternal UTI, time-mated dams were transurethrally infected with 5e7 colony forming units (CFU) bacteria on embryonic day (E)13.5 and monitored daily for signs of preterm birth (e.g. vaginal bleeding, passage of fetal tissue or amniotic fluid) until E17.5 (**Figure 1A**). Roughly 40% of dams exhibited signs of preterm labor within 4 hours of infection with UTI89, a well-characterized cystitis *E. coli* strain, compared to 0% in mock-infected dams (**Figure 1B**). UTI89, isolated in 1992 from a non-pregnant individual(21), may not accurately represent currently circulating UPEC, or pregnancy or preterm birth-related strains. Furthermore, *E. coli* evolves rapidly and has a broad pangenome(22). Given this, we selected two novel UPEC isolates, SL2 and SL323, isolated from pregnant human urine in 2023. SL2 was isolated from a patient with asymptomatic bacteriuria (ASB) who delivered at term gestation, while SL323 was from a symptomatic UTI patient who delivered preterm at 28 weeks of gestation. Similar to UTI89, 50% of dams infected with SL323 displayed preterm birth symptoms within four hours of infection (**Figure 1B**). On the other hand, ∼80% of dams infected with SL2 exhibited signs of preterm birth, although onset was delayed until 1.5 days post-infection in some cases (**Figure 1B**). Initiation of preterm birth did not require viable bacteria, as UV-inactivated UTI89 was sufficient to induce preterm birth (**Figure 1C**). Moreover, group B *Streptococcus* (GBS), the most common Gram-positive bacterium associated with ASB in pregnancy(23), also induced preterm birth in a subset of dams confirming this phenotype was not specific to UPEC (**Figure 1C**). Interestingly, a genetic knockout line for Tamm-Horsfall protein, shown to be more susceptible to UPEC UTI(24) exhibited similar preterm birth incidence in this model (**Figure S1**).

**Figure 1.**
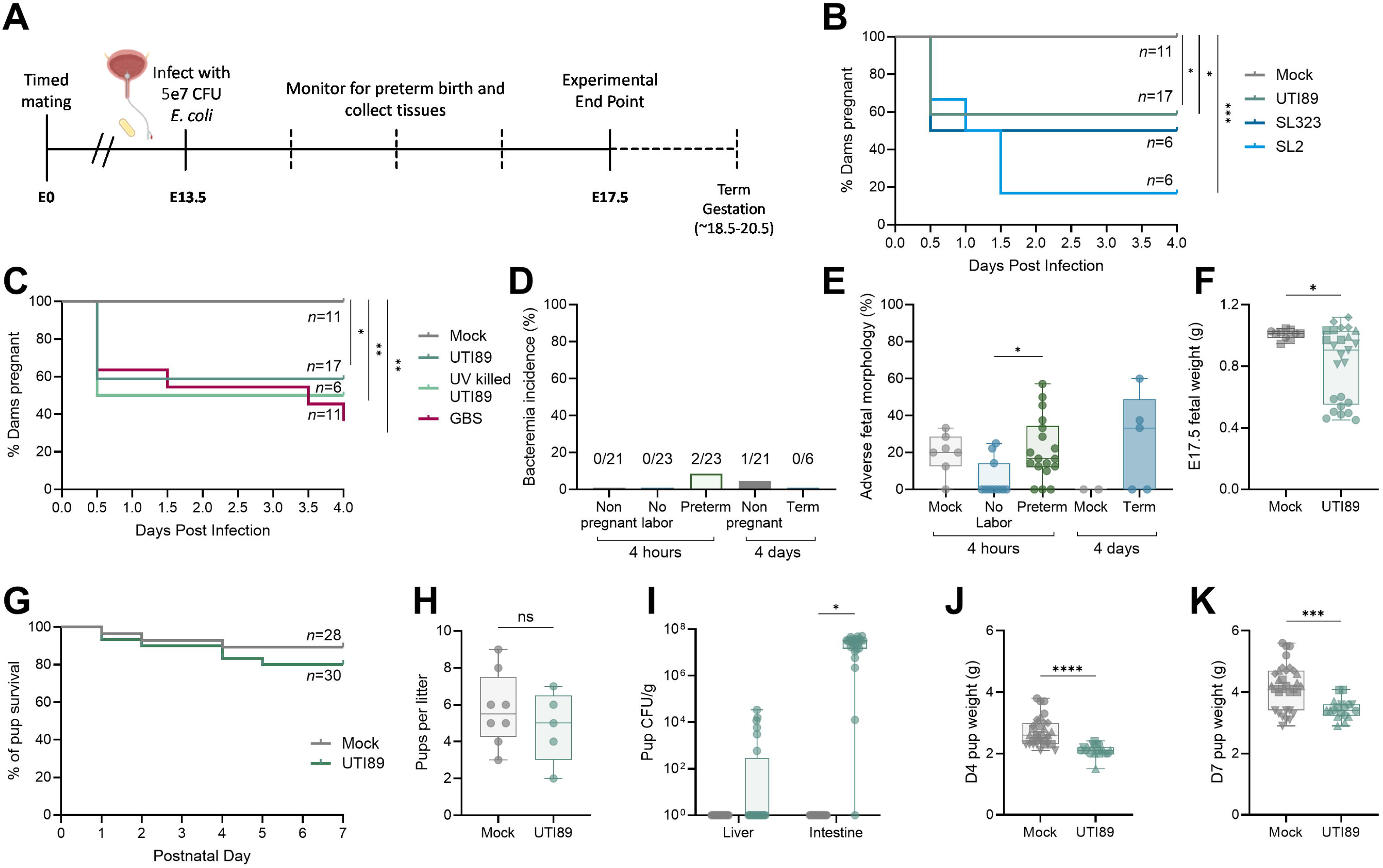
Murine maternal UTI mirrors clinical manifestations of UTI-associated adverse outcomes in human pregnancy. Dams were transurethrally infected with UPEC on embryonic day (E)13.5, then monitored for signs of preterm labor until E17.5 (**A**). Kaplan-Meyer curves depicting proportion of dams still pregnant over four days post-infection (**B,C**). Bacteremia incidence in pregnant and non-pregnant mice four hours and four days post infection (**D**). Adverse morphology proportions per litter for mock and infected dams four hours and four days post-infection (**E**). Fetal weight at E17.5 from mock or infected dams (**F**). Kaplan-Meyer curve depicting pup survival until post-natal day 7 from mock and infected dams (**G**). Litter size in mock and infected dams (**H**). Pup liver and intestinal bacterial burdens on postnatal day 7 from mock and infected dams (**I**). Postnatal day 4 (**J**) and 7 (**K**) weights from pups born to mock or infected dams. Experiments were performed at least twice with data combined. n=6-17 (B,C), n=6-23 (D), n=2-18 (E), n=10-25 (F), n=28-30 (G), n=5-8 (H), n=20-25 (I), n=19-32 (J), n=18-32 (K). Box and whisker plots show median, all points, and extend from 25^th^ to 75^th^ percentiles. Symbols are shaped by litter in F, J, K. Data were analyzed by Mantel-Cox test (B-C,G), Fisher’s exact test (D), Kruskal-Wallis test (E), Mann-Whitney (F, H, J-K), or two-way ANOVA with Benjamini, Krieger and Yekutieli correction for false discovery with a false discovery rate set at 5% (I). *p<0.05, **p<0.01, ***p<0.001, ****p<0.0001 (B-C, E-F, J-K). *q<0.05 (I).

Urosepsis is an increased and urgent complication of UTI in pregnancy(25). In this murine model, bacteremia following infection was rare and frequency did not significantly differ between groups (**Figure 1D**). Because the impact of systemic maternal inflammation on birth outcomes has been well-described(7), we sought to characterize the effects of localized urinary infection on maternofetal health; therefore, bacteremic dams were excluded from all subsequent analyses. Increased risk of neonatal complications, such as stillbirth, fetal growth restriction, and neonatal sepsis, are associated with maternal UTI(4, 10, 11). Using gross morphological assessment, we detected adverse fetal morphologies (reabsorption, fetal demise, fetal necrosis) more frequently in infected dams with preterm birth compared to non-laboring infected dams at four hours post-infection (**Figure 1E**). Although no differences in the rate of adverse morphologies was observed between infected dams and mock-infected controls at E17.5, fetal growth restriction was evident in the infected group (**Figure 1F**). To determine if there were postnatal impacts, we allowed dams to give birth and monitored pups out to postnatal day 7. There was no difference in pup survival (**Figure 1G**) or average litter size (**Figure 1H**). Most pups born to infected dams were intestinally colonized with UPEC, and a quarter of them (6/25) had UPEC disseminated to the liver (**Figure 1I**). Most dams exhibited urinary and reproductive tract colonization although total burdens varied across time points (**Figure S1**). Growth restriction in pups from infected dams was maintained at postnatal day 4 and 7 (**Figure 1J-K**). Overall, this model demonstrates multiple features of UTI-associated pregnancy complications in humans, including sporadic preterm birth and in utero growth restriction, in the absence of maternal systemic infection.

### Bladder inflammation during UTI is altered in pregnancy

There are several proposed reasons for increased risk for UTI in pregnancy, including anatomical and hormonal drivers of susceptibility(26). Pregnancy-associated systemic immunological changes are well-documented(7); however, the impact of pregnancy on bladder mucosal immunity remains unexplored. Using this murine model of maternal UTI, we examined the impact of pregnancy on the acute bacterial burdens and immune response to infection. Non-pregnant and pregnant mice (E13.5) were transurethrally infected with UTI89 and urinary and reproductive tract tissues were collected four hours post-infection. Bacterial burdens were similar in bladder and kidneys, and there were no differences in dissemination to the reproductive tract between pregnant and non-pregnant mice (**Figure 2A**). Additionally, we observed UPEC dissemination to the ileal lymph node, a shared draining lymph node between the urinary bladder and uterus(27), in a subset of pregnant dams that was not detected in non-pregnant mice (**Figure 2B**)(26). Bulk RNA sequencing and Reactome pathway analyses revealed 52 differentially regulated pathways, including an increase in MHC class II antigen presentation pathways in pregnant infected bladders compared to non-pregnant infected bladders, but a decrease in antigen cross presentation, antigen sensing (TLR4 and TLR9 cascades, C type lectin receptors), downstream TCR signaling, and signaling by interleukins (**Figure 2C, Supplemental Table 1**). Interestingly, we observed differential expression of several adhesins (*Ceacam14, Ceacam3, Gm5155*, and *Psg18*), known to be expressed in the placenta(28–30), but with unknown function in the bladder (**Figure 2D**). To determine if transcriptional differences translated to altered immune cell populations, we performed flow cytometry on whole bladders (determined according to gating scheme in **Figure S2**). In line with transcriptional profiling, we observed increased CD45+ proportions in nonpregnant bladders (**Figure S2**), marked by a significant decrease in dendritic cells (DC) in pregnant bladders during infection (**Figure 2E**). While IL-2 and IL-3 were elevated in the pregnant bladder, circulating RANTES was decreased in pregnant mice, supporting altered T cell transcriptional behavior (**Figure 2F-H**). Twenty other cytokines were measured with no significant differences (**Figure S3**). Together, these results indicate a general reduction in antigen sensing in the pregnant bladder during UTI.

**Figure 2.**
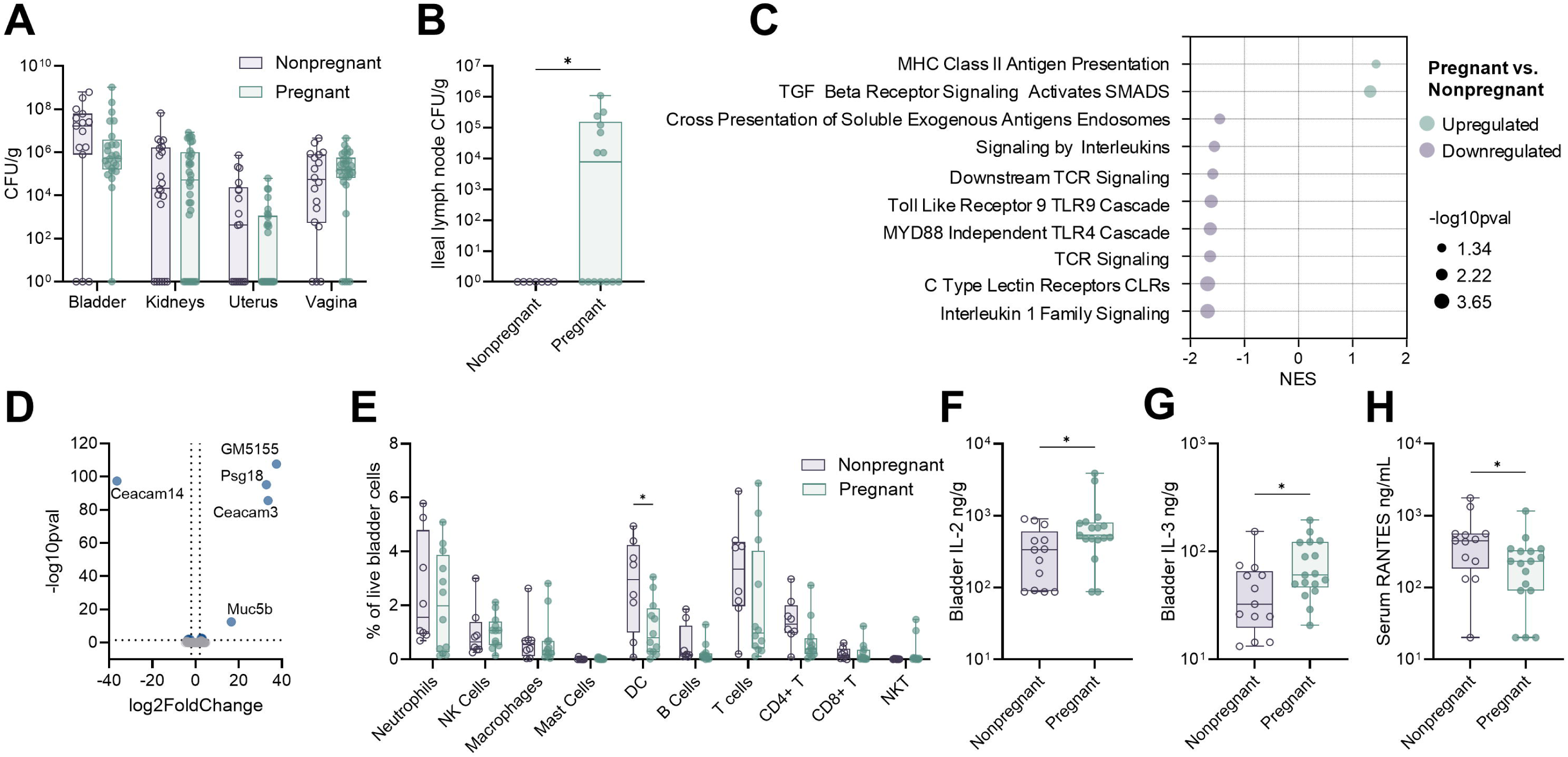
Bladder immunity to UTI is altered during pregnancy. Bladder, kidney, uterine/decidual, vaginal, and ileal lymph node bacterial burdens four hours post-infection in non-pregnant and pregnant mice (**A, B**). Reactome pathway enrichment (**C**) and differential gene expression (**D**) in pregnant bladders compared to non-pregnant bladders. Flow cytometry on pregnant and non-pregnant bladders four hours post-infection (**E**). Bladder IL-2 (**F**) and IL-3 (**G**) and serum RANTES (**H**) in infected pregnant and non-pregnant mice. Experiments were performed at least twice with data combined. n=15-33 (A), n=7-14 (B), n=4-11 (C-D), n=8-12 (E), n=13-17 (F-H). Box and whisker plots show median, all points, and extend from 25^th^ to 75^th^ percentiles. NES = normalized enrichment score. Data were analyzed by two-way ANOVA with Benjamini, Krieger and Yekutieli correction for false discovery with a false discovery rate set at 5% (A,E), fGSEA with a gene set minimum of 15 and maximum of 500 and 10,000 permutations using the Reactome collection from Molecular Signature Databases (C), generalized linear model, Log2 fold change >1 and Wald tests with FDR adjusted p valueL<L0.05 (D), or Mann-Whitney (F-H). *p<0.05 (F-H). *q<0.05 (A,E).

### Preterm dams experience increased acute inflammation in the bladder

Given the bimodal outcomes of either preterm birth or sustained pregnancy following bacterial inoculation, we interrogated whether infection dynamics or immune factors distinguished these groups. At four hours post-infection, urinary tract bacterial burdens were not different between dams exhibiting signs of preterm birth and those that were not (**Figure 3A**). Both infected groups demonstrated transcriptional upregulation of placenta-associated adhesins (*Ceacam3, Ceacam11, Gm5155, Psg18*) in bladders compared to mock pregnant controls as measured by bulk RNA sequencing, suggesting pregnancy-specific factors modify the immune response to infection at distal sites such as the bladder (**Figure 3B**). Transcriptional elevation of genes related to antigen sensing and neutrophil recruitment (*Clec4e, Cxcl2, Cxcr2, Il1b, Il36g*), interferon signaling (*Ifit1, Isg15, Oas3, Oasl1*), and alarmins (*S100a8, Saa3*) occurred in preterm dams compared to control dams, suggesting aberrant inflammation in the preterm bladder (**Figure 3B**). In line with this, there was significantly greater neutrophil and T cell infiltration in the bladder of preterm dams compared to mock and non-laboring groups (**Figure 3C**). Bladder neutrophils could be visualized in the uroepithelium and submucosa in both preterm and non-laboring dams (**Figure 3D**) but did not result in significant differences in histopathological scoring, likely due to the short timeframe following inoculation (**Figure 3E**). Multiplex cytokine analysis showed decreased IFN-γ and IL-3 in the bladders of preterm dams compared to non-laboring dams (**Figure 3F-H**) suggesting altered T cell activation. Consistent with our transcriptional results, S100a9 protein was not differentially detected between preterm and nonlaboring bladders but was significantly elevated compared to mock when infected groups were combined (**Figure 3I**). Together, these results suggest excessive inflammatory responses in the bladders of dams exhibiting preterm labor that are either absent or held in check in the subset of dams not undergoing preterm labor.

**Figure 3.**
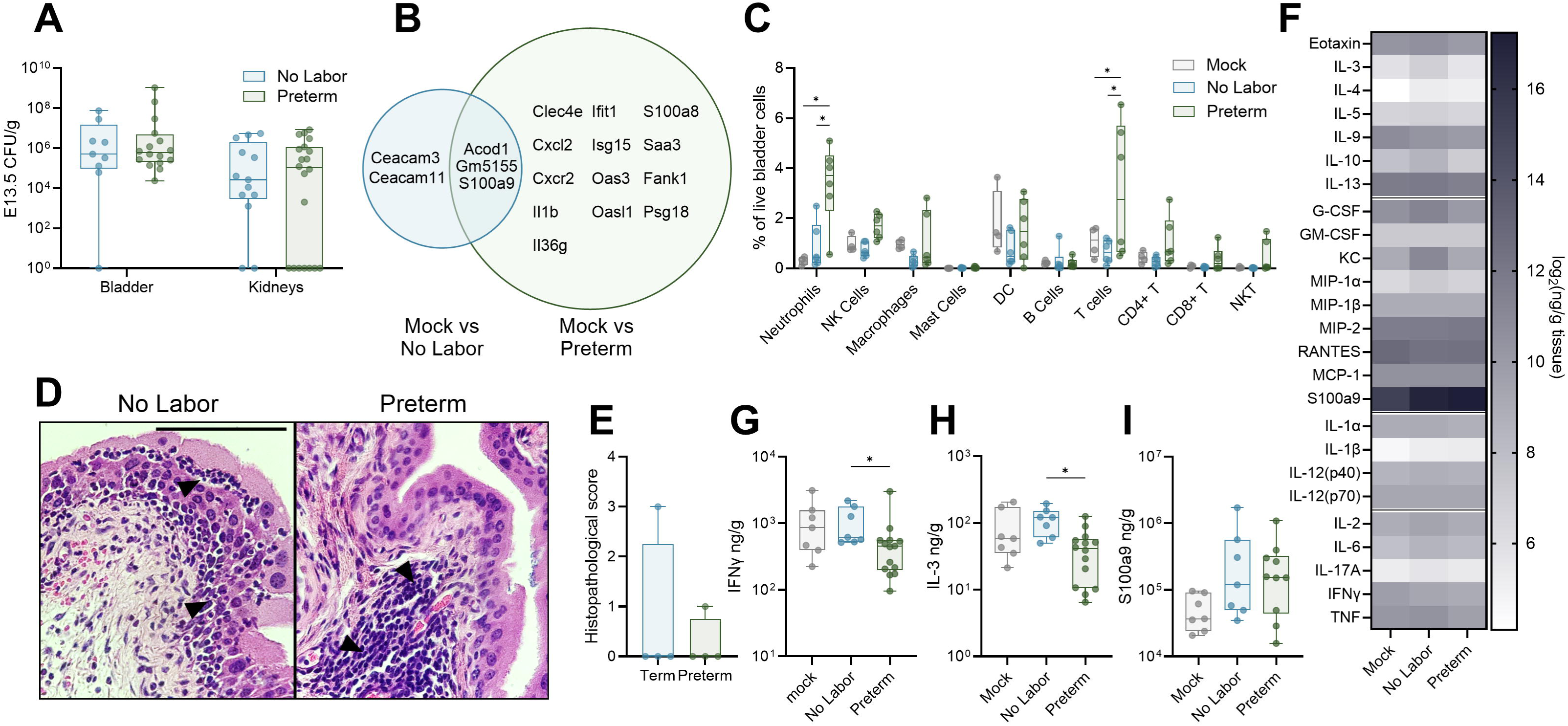
Preterm bladders display increased inflammation. Bladder and kidney bacterial burdens four hours post-infection (**A**). Venn diagram showing differentially expressed genes in preterm and non-laboring bladders compared to mock bladders (**B**). Flow cytometry on bladders of mock, non-laboring, and preterm dams four hours post-infection (**C**). Representative images (**D**) and histopathological scoring (**E**) of pregnant infected bladders. Black arrows indicate polymorphonuclear cell aggregates. Heatmap (**F**) and box and whisker plots (**G-H**) of select bladder cytokines in mock, non-laboring, and preterm dams. Experiments were performed at least twice with data combined. n=9-20 (A), n=3-6 (B), n=4-6 (C), n=4 (E), n=7-14 (F-H). Box and whisker plots show median, all points, and extend from 25^th^ to 75^th^ percentiles. Data in A and B represent pregnant samples from Figure 2A and 2E, respectively, disaggregated by outcome. Data were analyzed by two-way ANOVA with Benjamini, Krieger and Yekutieli correction for false discovery with a false discovery rate set at 5% (A,C), generalized linear model, Log2 fold change >1 and Wald tests with FDR adjusted p valueL<L0.05 (B), or Mann-Whitney (E,G-H). *p<0.05 (G-H). *q<0.05 (C).

### Maternal bladder infection induces uteroplacental inflammation

Maternal UTI is associated with various adverse pregnancy and neonatal outcomes, including preterm birth, preeclampsia, fetal growth restriction, and neonatal sepsis, often with the same organism isolated from maternal urine(23, 31–34), indicating localized bladder infection has distal impacts on the maternofetal interface. To characterize bacterial dissemination and reproductive immune responses, we collected maternal vaginal and decidual samples as well as placentae at four hours post-infection. UPEC was detected in the vaginal tract in most dams, but only ascended to the decidua in a third of mice (13/35) with no differences in burdens between preterm and non-laboring dams (**Figure 4A**). Bacterial dissemination to the placentae was quite rare at this timepoint with UPEC detected in only 11% (20/182) and no differences between groups (**Figure 4B**). At four days post-infection, placental dissemination was more common rising to a rate of 39% (13/33) (**Figure 4C**). No overt histopathology was observed in placentae from either preterm or non-laboring dams at four hours post-infection (**Figure 4D**). Even so, bulk RNA sequencing detected complement factor D (*Cfd*), a regulatory factor in the alternative complement pathway that has been associated with pre-eclampsia(35), as significantly elevated in preterm placentae (**Figure 4E**). Reactome pathway analyses revealed increased decidual antigen recognition, increased placental interleukin signaling, depressed B cell signaling, and disrupted hemostasis (platelet pathways and VEGF signaling) in preterm dams (**Figure 4F**). Flow cytometry revealed a shift in placental immune cell populations in infected mice, where non-laboring dams had greater placental neutrophil infiltration and preterm dams had increased placental T cells (**Figures 4G**). Decidual NK cells were elevated in the preterm group (**Figure S2**). Given the complex role T cells play at the maternal-fetal interface, we further characterized T helper subtypes in the placenta using intracellular transcription factor staining (determined according to gating scheme in **Figure S2**). Placentae from mock-infected dams showed greater proportions of naïve T_H0_ cells, whereas placentae from preterm dams and non-laboring dams showed increased T_H17_ polarization and FoxP3+RORγt+ T cell intermediates respectively (**Figure 4H**). Interestingly, there was increased G-CSF and MCP-1 in the preterm deciduae and placentae, suggesting initiation of recruitment of additional immune cells, such as monocytes, macrophages, and neutrophils (**Figure 4I-L**).

**Figure 4.**
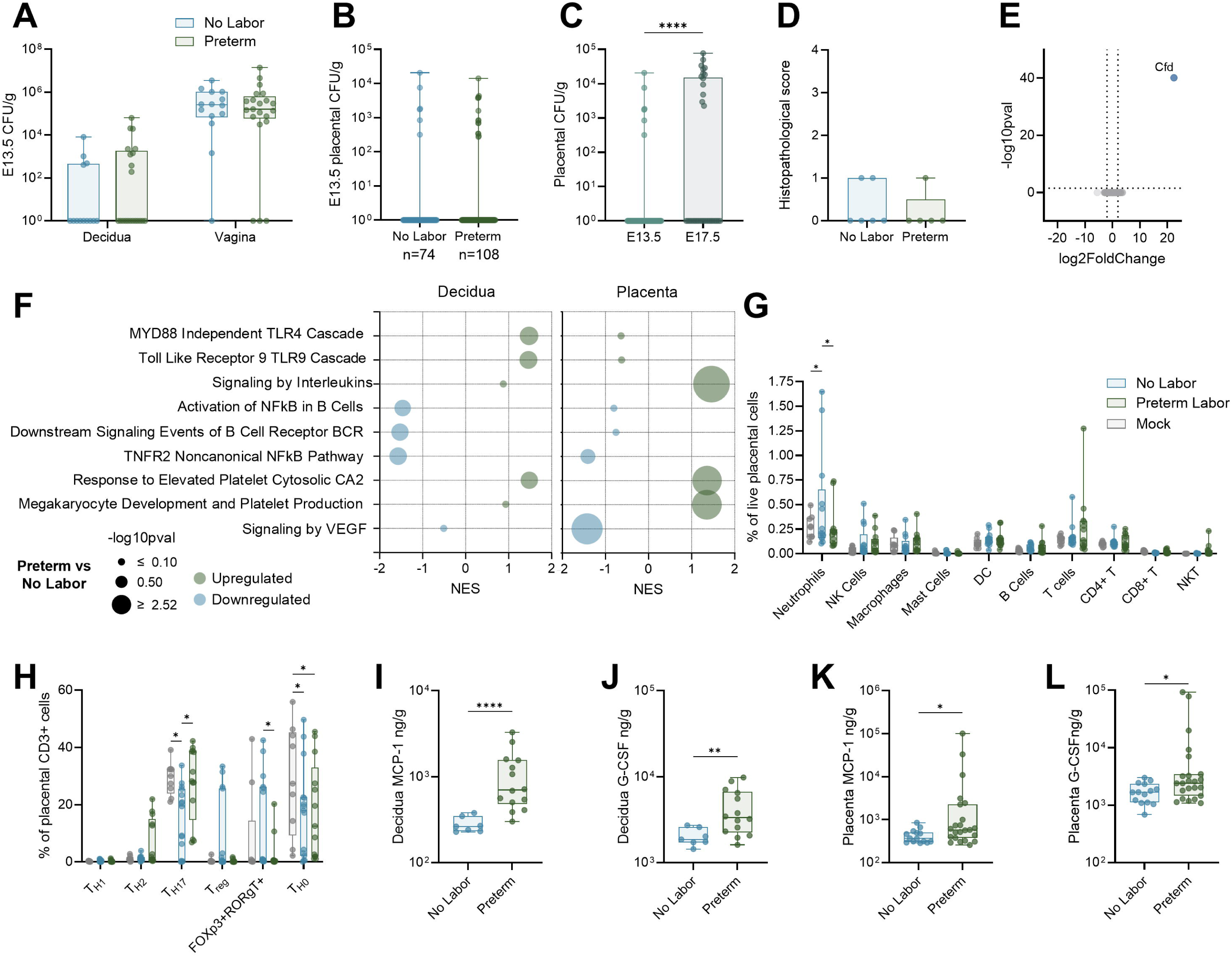
Uteroplacental immune activation is elevated in preterm dams. Decidual and vaginal bacterial burdens four hours post-infection (**A**). Placental bacterial burdens on E13.5 in nonlaboring and preterm dams (**B**) or on E13.5 and E17.5 in non-laboring dams (**C**). Placental histopathological scoring (**D**). Differentially expressed genes (**E**) and Reactome pathways (**F**) in preterm placentae (**E-F**) or deciduae (**F**) compared to non-laboring dams. Broad immunophenotyping (**G**) or T helper lymphocyte phenotyping (**H**) by flow cytometry of mock, non-laboring, and preterm placentae 4 hours post-infection. Decidual (**I-J**) and placental (**K-L**) MCP-1 and G-CSF cytokine levels in non-laboring and preterm dams four hours post-infection. n=14-21 (A), n=74-108 (B), n=74-33 (C), n=5-6 (D), n=5-12 (E-F), n=9-13 (G), n=9-15 (H), n=7-14 (I-J), n=14-23 (K-L). Box and whisker plots show median, all points, and extend from 25^th^ to 75^th^ percentiles. NES = normalized enrichment score. Data were analyzed by two-way ANOVA with Benjamini, Krieger and Yekutieli correction for false discovery with a false discovery rate set at 5% (A,G-H), Mann-Whitney (B-D, I-L), generalized linear model, Log2 fold change >1 and Wald tests with FDR adjusted p valueL<L0.05 (E), or fGSEA with a gene set minimum of 15 and maximum of 500 and 10,000 permutations using the Reactome collection from Molecular Signature Databases (F). *p<0.05, **p<0.01, ****p<0.0001 (C, I-L). *q<0.05 (G-H).

### Male sex is associated with increased placental inflammation and risk of preterm birth

Male fetal sex has been associated with increased risk of adverse perinatal and neonatal outcomes(1, 36), including increased risk of preterm birth. Preterm dams had a higher proportion of males compared to non-laboring dams as determined by PCR (**Figure 5A**), suggesting male fetuses may contribute to preterm birth risk in this litter-bearing model. The relationship between fetal sex and adverse outcomes, however, was complex. No sex differences in placental UPEC invasion or fetal growth restriction *in utero* were detected (**Figure 5B-C**), although males were less likely to experience stunted postnatal weight gain (**Figure 5D**). Flow cytometry also showed elevated T cells in the preterm male placentae (**Figure 5E**), although increased T_H17_ cells were detected in both male and female preterm placentae (**Figure 5F**). Elevated T_regs_ and FoxP3+RORγt+ T cell intermediates were more notable in female fetuses from nonlaboring dams(**Figure 5F**). Heightened inflammatory cytokines were seen in male fetuses, including increased placental G-CSF, MIP-1α, and MCP-1 in preterm males (**Figure 5G**). These findings suggest that fetal sex determines the extent of placental responses to maternal infection and contributes to infection outcome.

**Figure 5.**
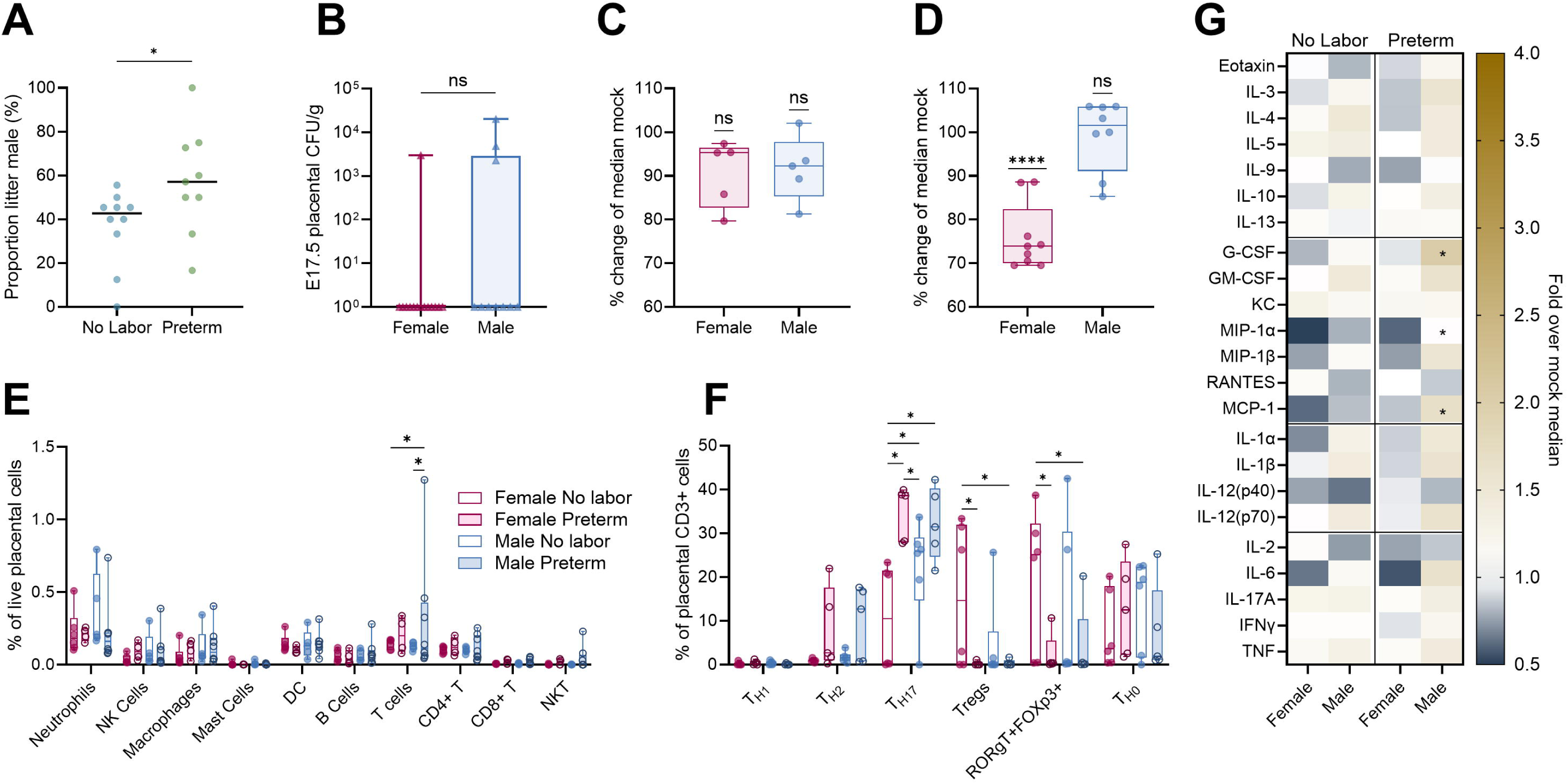
Fetal sex contributes to differential placental immunity and adverse outcomes. Male proportion of each litter in non-laboring and preterm dams (**A**). Placental bacterial burdens in males and females four days post-infection (**B**). Fetal E17.5 (**C**) and pup postnatal day 7 weight (**D**) from infected dams shown as a proportion of median fetal or pup weight of the same sex from mock-infected dams. Broad immunophenotyping (**E**) and T helper lymphocyte immunophenotyping (**F**) of placentae stratified by fetal sex and maternal outcomes. Placental cytokine quantification stratified by fetal sex and maternal outcome displayed as fold change over median placentae of the same sex from mock-infected dams (**G**). Experiments were performed at least twice with data combined. Data in C, D, E, and F represent data from Figure 1F, Figure 1K, Figure 4G, Figure 4H, respectively, disaggregated by sex. n=9-10 (A), n=10-14 (B), n=5 (C), n=8-9 (D), n=4-8 (E), n=5-6 (F), n=7-10 (G). Box and whisker plots show median, all points, and extend from 25^th^ to 75^th^ percentiles. Data were analyzed by Mann-Whitney (A-B), one sample T test against the theoretical value of 100% (C-D), or two-way ANOVA with Benjamini, Krieger and Yekutieli correction for false discovery with a false discovery rate set at 5% (E-G). *p<0.05, ****p<0.0001 (A, D). *q<0.05 (E-G).

### Elevated systemic IL-10 protects from preterm birth

Next, we examined if the maternal systemic immune response differed between preterm and non-laboring dams via cytokine multiplex array. A significant increase in circulating IL-10 was the only differentially detected cytokine in the non-laboring group, suggesting a protective regulatory response that maintained pregnancy in those dams, though no changes in splenic immune cells were observed (**Figure 6A, Figure S2, Figure S3**). To test the causative role of maternal systemic IL-10, we treated dams with recombinant IL-10 via intraperitoneal injection immediately after infection (**Figure 6B**). While 41% of untreated dams were preterm (7/17), only 11% (1/9) were preterm in the IL-10 group (**Figure 6C**). Although incidence of preterm birth was reduced, IL-10 treatment also corresponded with intrauterine growth restriction in both infected and mock-infected groups (**Figure 6D**). Additionally, even though IL-10 exerts immunosuppressive effects, we observed no difference in maternal urinary or reproductive tract bacterial burdens or placental invasion (**Figure 6E-F**). In non-pregnant females, there was a slight, but significant, reduction in kidney bacterial burdens (**Figure 6E**). These data suggest that immunoregulation during infection contributes to pregnancy maintenance, but other aspects of fetal health may be impacted.

**Figure 6.**
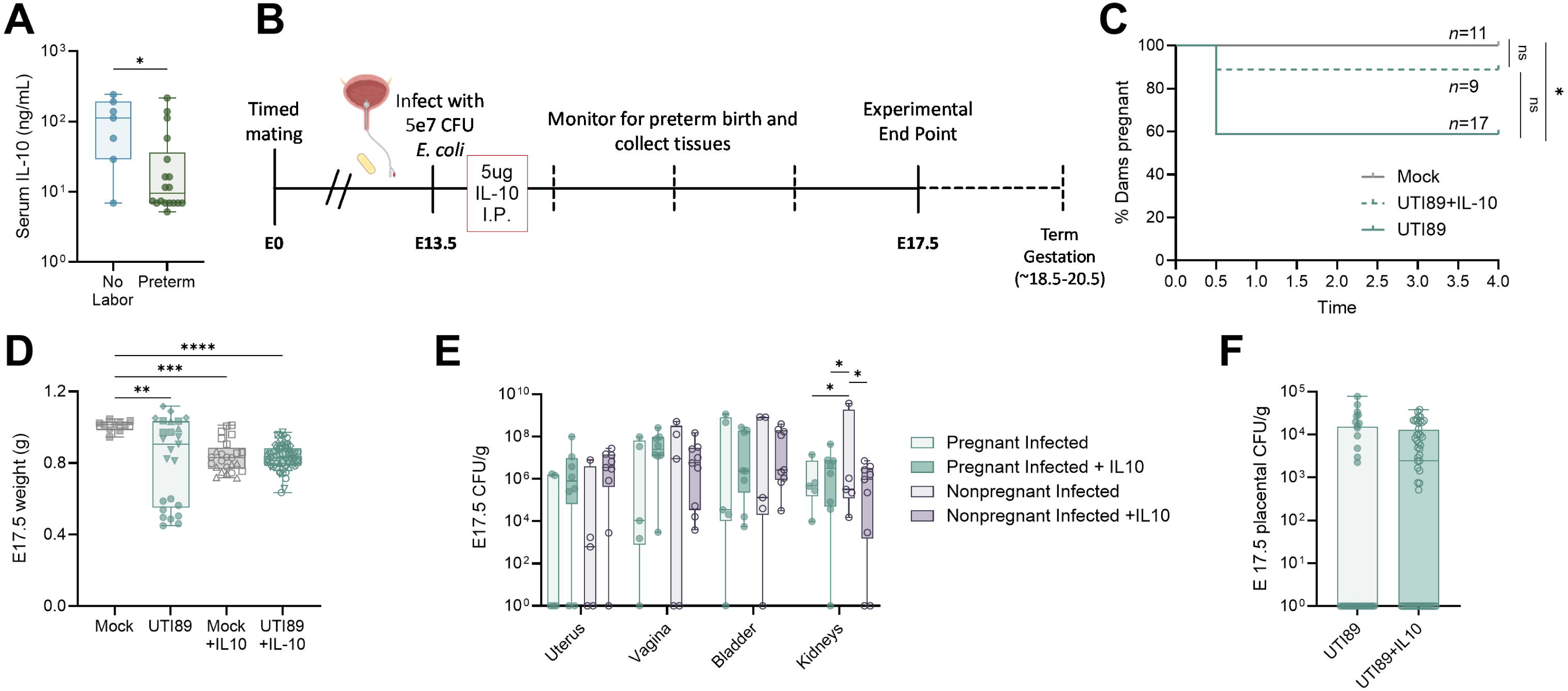
IL-10 is lower in preterm birth and associated with fetal growth restriction. IL-10 quantified in maternal serum (**A**). Immediately after infection, mice were treated with 5µg IL-10 in PBS by intraperitoneal injection (**B**). Kaplan-Meyer curve depicting rate of ongoing pregnancy in dams in days post-infection (**C**). E17.5 fetal weight (**D**). Uterine, vaginal, bladder, and kidney bacterial burdens in pregnant and non-pregnant mice with and without IL-10 treatment (**E**). E17.5 placental bacterial burdens with and without IL-10 treatment (**F**). Experiments were performed at least twice with data combined. n=7-18 (A), n=9-17 (C), n=10-54 (D), n=5-9 (E), n=33-52 (F). Box and whisker plots show median, all points, and extend from 25^th^ to 75^th^ percentiles. Data were analyzed by Mann-Whitney (A,F), Mantel-Cox test (C), Kruskal-Wallis test (D), or two-way ANOVA with Benjamini, Krieger and Yekutieli correction for false discovery with a false discovery rate set at 5% (E). *p<0.05, **p<0.01, ***p<0.001, ****p<0.0001 (A,D). *q<0.05 (E).

### Urinary cytokines differ between UTI and ASB and throughout gestation in humans

To evaluate maternal immune responses to UTI and ASB in pregnancy, we performed a cross-sectional cohort study to evaluate urinary cytokines in pregnant and non-pregnant individuals. Our cohort consisted of 59 urine samples from singleton pregnancies (22 control, 18 ASB, 19 UTI) and 6 urine samples from non-pregnant, reproductive age females (8 control, 6 UTI) (**Supplemental Table 2**). Samples were collected throughout pregnancy, ranging from 5.0 to 39.7 weeks in gestation. Birth outcomes, including gestational age at birth, fetal sex, and fetal weight at birth are known for 81% of the cohort (48/59). Preterm birth was high in this cohort at 38% (18/47) of live births (**Supplemental Table 3**). Using a multiplex cytokine array, we detected elevated IL-8 and GROα in pregnant UTI patients compared to control and ASB patients (**Figure 7A**). No cytokines were significantly upregulated in ASB samples compared to controls, suggesting limited urinary inflammation in ASB patients (**Figure 7A**). Multiple urine cytokine levels correlated with gestational age. A total of 16 cytokines were significantly correlated with gestational age in culture positive urine samples (ASB and UTI) at sample collection by linear regression (**Figure S4**), only 5 of which were correlated with gestational age in negative culture samples (**Figure S5**). Specifically, four cytokines (IL-9, IL-22, IL-17F, and GM-CSF) are associated with T_H17_ differentiation and activation (**Figure 7B-E**). When stratifying culture negative samples by fetal sex, higher IL-1α and IL-8 were detected in urine from pregnancies with female fetuses, while IL-5 was higher in urine from pregnancies with male fetuses (**Figure 7F-H**).

**Figure 7.**
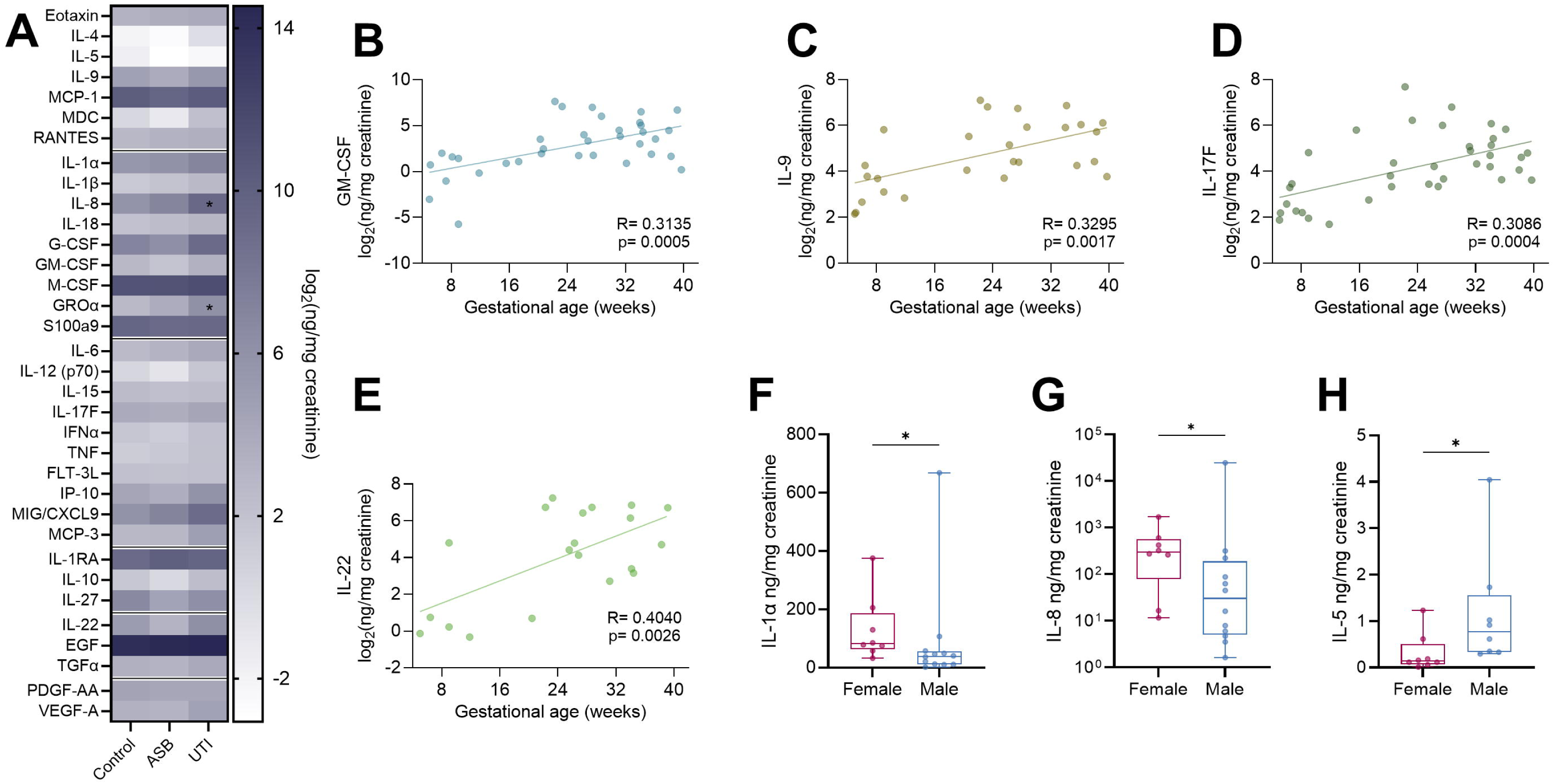
Gestational age and fetal sex correlate with urine cytokines. Heat map comparing urinary cytokines from pregnant patients with asymptomatic bacteriuria (ASB), diagnosed urinary tract infection (UTI), or culture negative controls (**A**). Correlation of the log_2_ transformation of urinary cytokines normalized to creatinine with gestational age at sample collection in patients with positive urine culture (**B-E**). Urinary cytokines normalized to creatinine stratified be fetal sex in bacterial culture negative samples (**F-H**). For all panels, samples that fell below the limit of detection were excluded. n=18-22 (A), n=35 (B), n=27 (C), n=37 (D), n=20 (E), n=8-12 (F,G), n=8 (H). Box and whisker plots show median, all points, and extend from 25^th^ to 75^th^ percentiles. Data were analyzed by two-way ANOVA with Benjamini, Krieger and Yekutieli correction for false discovery with a false discovery rate set at 5% (A), simple linear regression (B-E), or Mann-Whitney (F-H). *p<0.05.

### Urinary cytokines correlate with preterm birth with and without stratifying by urine culture status

We performed Spearman correlation analyses of the 34 detected urinary cytokines with gestational age at birth, gestational age a sample collection, and birth weight. As expected, gestational age at birth and fetal weight were positively correlated in all groups (**Figure 8A**). In negative urine culture samples, ten cytokines (RANTES, IL-1β, IL-18, M-CSF, IL-15, L-17F, IFNα, TNF, FLT-3L, and EGF) were significantly inversely correlated with gestational age at delivery, while IL-18 and IL-15 were correlated with gestational age at sample collection (**Figure 8A**). MCP-1, RANTES, IL-1β, S100A9, IL-15, and IL-10 were all inversely correlated with birth weight in the negative culture group (**Figure 8A**). These data suggest elevated cytokines in the urine in the absence of bacteriuria is associated with poorer pregnancy outcomes. Using these data, we developed a four-cytokine model to predict preterm birth in the absence or presence of bacteriuria, or with all samples combined. IL-1β and EGF were significantly higher in the culture negative preterm group (**Figure 8B-C**). While TNF and IL-17F were not significantly increased, adding them to the multiple linear regression model increased the sensitivity and specificity of the model with an AUC of 0.9 (**Figure 8D-F**). In the positive urine culture group, FLT-3L and PDGF-AA were significantly correlated with gestational age at delivery (**Figure 8A**). When combined with EGF and RANTES, we generated a linear regression model that predicted preterm birth risk with an AUC of 0.9063 (**Figure 8G-K**). When we analyzed the combined cohort, S100a9 and IL-10 remained significantly inversely correlated with birth weight (**Figure 8A**). When we grouped samples as above or below the median sample value, samples with above median IL-10 resulted in significantly lower birth weights (**Figure 8L**). While no cytokines were significantly different between term and preterm cases in the combined group, incorporating IL-10, IL-1RA, IL-15, and IL-1β into a multiple linear regression model yielded an AUC of 0.8235 and significantly predicted preterm birth (**Figure 8M-Q**). Together, these data reveal a potential prognostic value of urine cytokine levels and birth outcomes in a human cohort.

**Figure 8.**
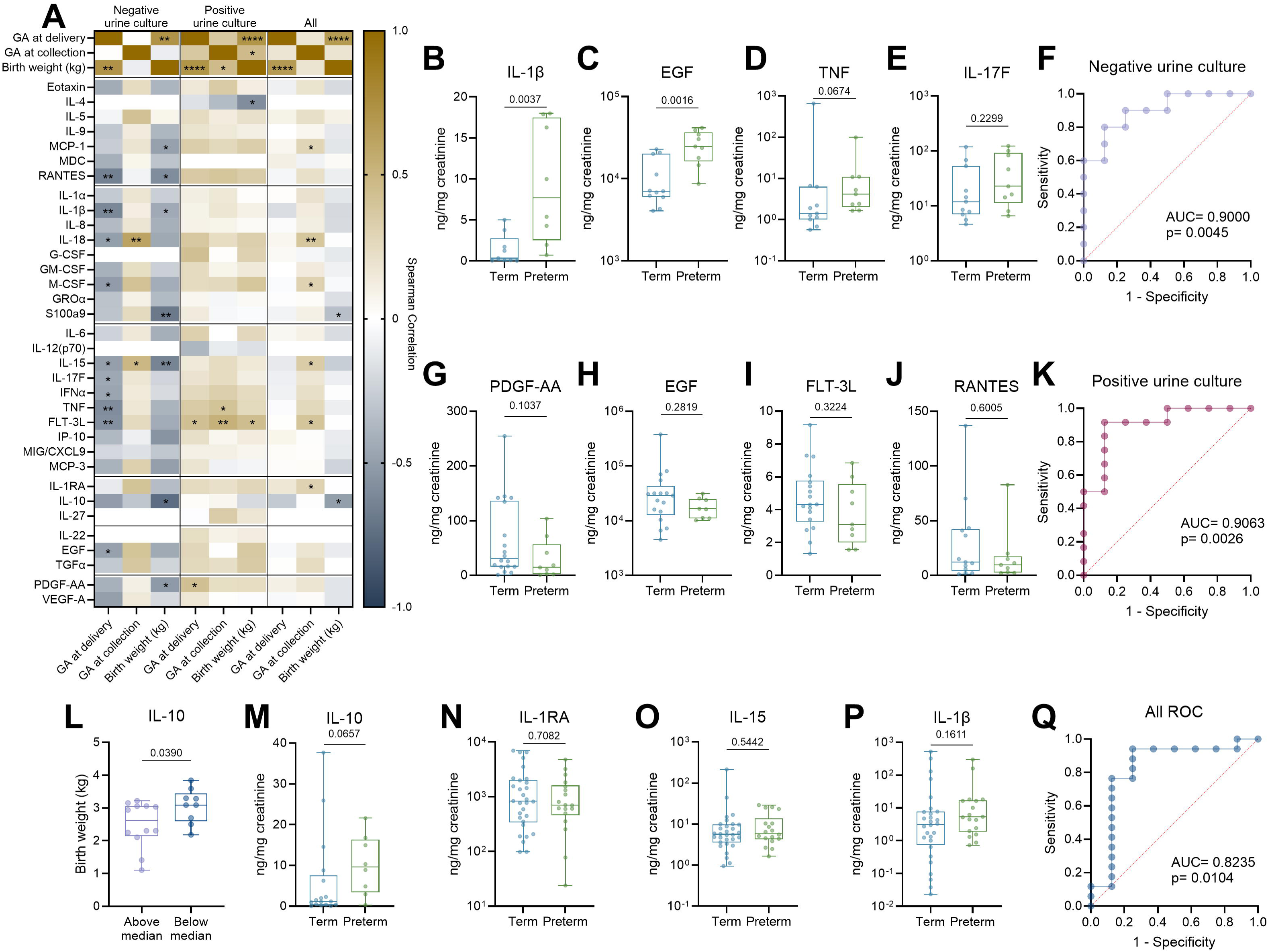
Urinary cytokines predict preterm birth risk. Heatmap of Spearman correlations between log_2_ transformation of 34 urinary cytokines normalized to creatinine and gestational age (GA) at sample collection, GA at birth, and birth weight (**A**). Urinary cytokine quantification stratified by birth outcome in culture negative samples (**B-E**). Receiver-operating characteristic (ROC) curve for a four-cytokine model to predict preterm birth risk in culture negative urine (**F**). Urinary cytokine quantification stratified by birth outcome in culture positive samples (**G-I**). ROC curve for a three-cytokine model to predict preterm birth risk in culture positive urine (**J**). Birth weight of neonates born to patients in this trial stratified as above or below the median cytokine concentration (**K,L**). Urinary cytokine quantification stratified by birth outcome in all samples regardless of culture (**M,N,O,P**). ROC curve for a four-cytokine model to predict preterm birth risk in all samples regardless of culture (**Q**). For all panels, samples that fell below the limit of detection were excluded. n=17-40 (A), n=7-8 (B), n=8-9 (C-E), n=17 (F), n=7-15 (G), n=6-14 (H), n=7-16 (I), n=23 (J), n=8-15 (K), n=16-24 (L,M,N), n=40 (O), n=8-12 (P), n=13-22 (Q). Box and whisker plots show median, all points, and extend from 25^th^ to 75^th^ percentiles. Data were analyzed by simple linear regression with spearman correlation (A), Mann-Whitney (B-E,G-I,K-O), or with multivariate logistic regression with test of significance for area under the ROC curve (F,J,Q).

## DISCUSSION

Clinical data have solidified the significance of UTI in pregnancy, establishing increased prevalence and multifactorial peri- and postnatal consequences. Despite striking correlations, mechanistic data investigating immunological and bacterial drivers of UTI-associated adverse pregnancy outcomes is limited. Here, we describe a murine model of maternal UTI which recapitulates clinical manifestations of maternal UTI observed in patients including preterm birth and intrauterine growth restriction. Preterm birth does not require bacterial dissemination to the reproductive tract in this model, suggesting maternal immune response initiates adverse outcomes. Multiple bacterial strains and species initiated preterm birth, albeit with different kinetics; thus, bacterial drivers of disease severity cannot be discounted. A retrospective cohort study showed variable preterm birth incidence depending on uropathogen; further, they found disease severity was associated with increased preterm birth incidence(3). Bacterial virulence factors, such as alpha hemolysin, cytotoxic necrotizing factor 1, and aerobactin, are associated with increased disease severity in UPEC UTI(37). Untangling the complex interplay between bacterial virulence factors, bladder inflammation, and adverse reproductive outcomes related to UTI is essential to design of therapeutic intervention.

Systemic immunomodulation during pregnancy has been well-described, but the impact of pregnancy on bladder immunity is underexplored. Our model demonstrates altered mucosal immunity in the pregnant bladder, driven by dendritic cell populations and T lymphocyte signaling, consistent with prior literature showing suppressed serum and urinary IL-6 in pregnant pyelonephritis patients compared to non-pregnant patients(38). However, a limitation of this model is that only one time point of gestational age at infection was evaluated. While UTI in the third trimester is more strongly linked to preterm birth and neonatal sepsis, multiple clinical studies have shown UTI and ASB are most common in the first trimester(3, 39, 40). Consistent with these findings, our cross-sectional cohort study identified a bacteriuria-specific positive correlation with multiple cytokines and gestational age, including those associated with T_H17_ function. Whether these urinary tract responses are suppressed in first trimester, or augmented in late gestation, remains unknown; however, we speculate several pregnancy-specific factors may contribute. Hormones, including human chorionic gonadotropin (hCG), estrogen, and progesterone, are detected in the urine(41, 42). Interestingly, hCG peaks in the first trimester and has been shown to induce T_regs_ and restrict T_H17_ cells(43). On the other hand, estrogen, which increases throughout gestation, may have a protective, immune enhancing role in the bladder by reducing UPEC, upregulating TNF, and downregulating IL-10(44, 45). To our knowledge, our study is the first to describe transcription of pregnancy specific glycoproteins (PSGs) in the bladder. Research on PSGs is limited; the placental-derived molecules have been detected in maternal circulation and may contribute to pregnancy maintenance including induction of IL-10(30, 46–48). It remains to be seen if alterations in mucosal immunity during pregnancy are attributed to systemic or local effects. Further, the impact of the gravid urinary environment on bacterial behavior remains uncharacterized.

In our model, increased inflammation was detected at the maternal-fetal interface despite limited bacterial dissemination to the uteroplacental space consistent with prior work(13) albeit with distinct differences. Bolton et al. observed increased polymorphonuclear cells (likely predominantly neutrophils) and macrophages in the uteroplacental space four days post-infection(13) whereas we observed increased neutrophils and T cells at four hours post-infection. Serum cytokines were elevated at four days post-infection and at term delivery in the Bolton model, while limited differences in circulating cytokines were observed in our model. These distinctions could both be attributed to differences in infection windows. Even so, they observed increases in serum IL-4, IL-17, IFN-γ, and IL-6, consistent with increased placental T_H2_ and T_H17_ populations in our model. An important outstanding question is the means by which bladder inflammation induces uteroplacental inflammation. Several studies have found correlations between systemic inflammation and distal site inflammation during UTI(13, 14); however, systemic cytokines and splenic immune populations were unchanged four hours post-infection in preterm and non-laboring dams compared to controls. Alternatively, bacterial dissemination from the urinary tract to the reproductive tract could induce inflammation; however, bacterial dissemination to the placenta was rare in our model and not detected by Bolton et al. Another possible mechanism of bladder-uteroplacental signaling is through lymphatic immune activation. The ileal lymph nodes drain the pelvic floor, including the bladder, uterus, and vaginal tract(27). We detected UPEC in the ileal lymph nodes in half of infected dams within four hours indicating immune cell trafficking from infected tissues. While during healthy pregnancy, fetal antigens are presented within lymph nodes to induce regulatory T cells(9), the impact of bacterial lymph node dissemination on maternofetal tolerance remains to be determined. The rapid onset of preterm labor in our model suggests that T cells in the placenta are likely polarized memory T cells rather than naïve T cells, which were reduced in placentae of infected dams. Effector memory and central memory T cells accumulate in the decidua during healthy pregnancy(49). Greater proportions of central memory cells in circulation have been described in pre-eclampsia and recurrent pregnancy loss, but their role in preterm birth is unknown(49). A caveat of this study is the use of a syngeneic pregnancy model which limits the capacity to evaluate maternal tolerance of foreign antigens. Despite this, T cell-mediated disruptions to pregnancy maintenance were still observed and the bimodal phenotype enabled us to interrogate differential immune responses that a more pronounced model may limit.

Male fetal sex is consistently linked to worse perinatal outcomes, including increased risk of preterm birth, pre-eclampsia, and gestational diabetes(1, 36). Male neonates also exhibit lower one-minute Apgar scores and increased odds of severe morbidity when preterm(19, 20). While most human pregnancies are singletons, mice gestate litters, which limits our ability to interrogate the impact of fetal sex on pregnancy outcomes. As a surrogate, we evaluated the proportion of males in each litter and found a greater proportion of males in preterm litters. The relationship between fetal sex and adverse outcomes is poorly defined but may be attributed in part to sex-specific maternal immunological adaptations. Female fetal sex has been associated with increased peripheral cytokines at baseline and in response to LPS(50, 51). In culture negative urine, we observed increased IL-1α and IL-8 in pregnancies with female fetuses, while IL-5 was increased in pregnancies with male fetuses. To our knowledge, this is the first study to characterize the impact of fetal sex on maternal urine cytokines. Some studies have linked sex-dependent factors with maternal bacteriuria, demonstrating placental TNF, IL-1β, and IL-10 in females negatively correlated with maternal bacteriuria(52, 53). It is unclear, though, if the relationship between fetoplacental inflammation and maternal bacteriuria is causative or correlative. The local uteroplacental immune microenvironment is also sexually dimorphic. Male placentae are marked by increased T_reg_ expansion(54). We did not see sex differences in placental T_regs_ in our model, potentially due to the syngeneic pregnancy. Interestingly, we found macrophage-associated cytokines were higher in preterm male placentae than preterm female in line with a prior study demonstrating male-specific elevated placental cytokines upon maternal immune activation with poly I:C(55). Sexually dimorphic behavior in placental macrophages (Hofbauer cells) has been recently described(17, 56). We observed no differences in placental macrophages quantities based on fetal sex in this study; however, we employed a limited characterization of macrophage phenotype. Sex-based differences may emerge with further discrimination between maternal and fetal macrophages or proinflammatory and anti-inflammatory macrophages.

The role of IL-10 in pregnancy is complex. IL-10 is a critical protective factor against LPS-induced pregnancy loss and preterm birth(57–59). In humans, serum IL-10 concentrations below the limit of detection are predictive of preterm birth risk(60, 61), reinforcing our finding that IL-10 plays a role in pregnancy maintenance during maternal UTI. While IL-10 promotes pregnancy maintenance, it may also augment infection susceptibility during pregnancy in some cases(62), but this was not observed in our model. Intrauterine growth restriction (IUGR) was exacerbated by exogenous IL-10 in both infected and mock-infected dams; however, this could be attributed to the high dose (∼30X the median detected concentration in infected mice) as a lower IL-10 dose (2.5µg vs 5µg) rescued LPS-induced growth restriction(59). Although other factors may contribute to IUGR, several additional observations support the link between IL-10 and IUGR: maternal urinary IL-10 inversely correlated with birth weight in our human cohort, and Bolton et al. reported concurrent increased circulating IL-10 and IUGR in their experimental UTI model(13). IL-10 is produced by both innate and adaptive immune cells, including monocytes, macrophages, dendritic cells, and B and T lymphocytes(63). T_regs_ and T_H2_ cells in the placenta are potential sources of IL-10, though IL-10 was not elevated in the decidua or placenta in our model. Further, there were no changes in helper T cell phenotypes in the maternal spleen. Regulatory B cells have been identified as important regulators of T cell polarization, particularly for their production of IL-10 and protection from inflammation-induced fetal demise(58). While we detected no changes in B cell populations in the placentae or spleen, B cell activation and signaling pathways were transcriptionally downregulated in the placentae of preterm dams.

Preterm birth is a multifarious condition with numerous etiologies, making it hard to predict and prevent(2). Cervical length is a reliable predictor of preterm birth risk in patients with cervical insufficiency but has limited efficacy in predicting spontaneous preterm birth(2, 64). Recent efforts have sought to identify biomarkers in maternal or fetal tissues(65). Elevated inflammatory markers, such as IL-1β, neutrophil elastase, and IL-8, in amniotic fluid are potential biomarkers for preterm birth(66–68); however, the invasive nature of amniocentesis drives increased risk of pregnancy loss and is typically not performed without prior indication(69). Cervicovaginal fluid provides a promising non-invasive alternative, and multiple studies report increased cervicovaginal IL-1β in spontaneous preterm birth samples(70, 71). Likewise, we found IL-1β was elevated culture negative urine from pregnancies that resulted in preterm birth. These findings underscore the therapeutic potential of IL-1 pathway blockade(72–75). Other studies have associated urinary oxidative stress markers, eicosanoids, and phthalates with preterm birth risk(76–78), but inflammatory markers are less characterized, although clinical studies are ongoing(79). Here, we propose three novel models employing urinary cytokines with high predictive value of preterm birth risk in culture negative, culture positive, and culture agnostic samples. The three-model system enables evaluation of preterm birth risk in a general population, but also facilitates specialized evaluation of culture positive samples when infection may confound basal urinary inflammation. Irrespective of culture status or gestational age, levels of IL-1RA, IL-10, IL-15, and IL-1β predicted preterm birth. As discussed above, IL-10 and IL-1 pathways have been implicated in preterm birth. IL-15 is an abundant uteroplacental cytokine that is increased in the amniotic fluid of preterm birth cases(80) and both augmented and deficient IL-15 levels are associated with poor pregnancy outcomes in vivo(81). Maternal factors can contribute to preterm birth risk, including factors such as race, body mass index, and maternal age(2, 82). Given the size of our cohort, we are unable to evaluate the contribution of maternal factors in our model. Preterm birth risk has been shown to be related to gestational age at UTI onset(3). Our cohort represents samples collected all throughout gestation but is underpowered to make trimester-specific predictions. An expanded study may illuminate more pronounced markers when gestational age at sample collection is considered.

In summary, our murine model of UTI-associated preterm birth provides a powerful tool to further investigate the molecular constituents of localized extrauterine infection that dictate perinatal and neonatal outcomes. We complemented in vivo findings with a human cohort, culminating in a novel, non-invasive model to predict preterm birth risk via urine cytokine analyses. Taken together, this preclinical model combined with easily accessible patient samples provides a unique platform to identify diagnostic and therapeutic opportunities in maternal-fetal health.

## MATERIALS AND METHODS

### Bacterial strains and culture conditions

*E. coli* strains UTI89(21), SL2 (UPEC isolate from a human pregnant ASB urine sample collected as a part of this study), and SL323 (UPEC isolate from a human pregnant UTI urine sample collected as a part of this study) were grown to stationary phase at 37°C with aeration in Luria broth (LB, Hardy Diagnostics) for at least 16 hours. *Streptococcus agalactiae* strain COH1 (ATCC, BAA-1176) was grown to stationary phase at 37°C in Todd-Hewitt broth (Hardy Diagnostics) for at least 16 hours. Bacterial cultures were centrifuged (3,220 × g for 5 minutes), washed with sterile PBS, then resuspended in the same volume of sterile PBS. For some experiments, overnight cultures were washed and resuspended in sterile PBS, then exposed to UV radiation for 1 hour. UV-inactivation was confirmed by plating on LB agar.

### Mice handling and breeding

Wild type female and male 129/sv mice aged 6 to 12 weeks were housed at BCM(83). Male mice were used exclusively for mating, while female mice were used for all in vivo experiments. Treatment and control groups were assigned randomly and housed 4 mice per cage. Mice ate and drank ad libitum and had a 12-hour light cycle. Timed mating was performed by co-housing one male with three females overnight. Soiled male bedding was added to female cages two days prior to mating to synchronously induce estrus and promote successful mating. Embryonic day 0.5 (E0.5) was considered as noon the day after mating. Mated females were weighed on embryonic days 0.5 and 10.5. Mice that gained at least 2 grams by E10.5 were considered likely pregnant. All animal protocols and procedures were approved by the BCM Institutional Animal Care and Use Committee under protocol AN-8233.

### In vivo model of urinary tract infection-associated preterm birth

An established mouse model of UTI was previously described(84, 85). On day 13.5 of gestation or 13.5 days post-coitus in non-pregnants (failed mating), mice were anesthetized via inhaled isoflurane, and approximately 5×10^7^ colony forming units (CFU) bacteria resuspended in sterile PBS were instilled transurethrally in 50 µL volume into the bladders. Mock-infected mice were treated likewise with 50 µL of sterile PBS. Mice were then monitored daily for vaginal bleeding, a sign of imminent labor. If vaginal bleeding was detected, mice were euthanized. If no vaginal bleeding was detected, mice were euthanized at 17.5 days of gestation. Upon euthanasia, whole blood was collected via cardiac puncture and plated on LB to assess bacteremia, then allowed to coagulate at room temperature for at least 30 minutes. After coagulation, clots were pelleted via centrifugation at 10,000 × g for 10 minutes and serum was removed and stored for later analysis. Bladder, kidneys, vaginal tract, and ileal lymph nodes were removed and homogenized in tubes containing sterile PBS and 1.0-mm-diameter zirconia/silica beads (Biospec Products; catalog number 11079110z) using a MagNA Lyser instrument (Roche Diagnostics). The uterine decidua was grossly dissected away from excess uterine tissue, and deciduae and placenta were homogenized as described above. Fetal gross morphology was assessed by two independent evaluators and categorized as healthy, reabsorbed, or fetal necrosis/demise. Fetal weight was recorded. Serial dilutions of bladder, kidney, placenta, and lymph nodes were plated on LB and bacteria was enumerated the next day. Serial dilutions of vaginal tract and decidua were plated on ChromAgar Orientation (ChromAgar, RT412), and purple, undulate colonies (indicating uropathogenic *E. coli*) were enumerated the next day.

To determine pup outcomes, dams were monitored twice daily for signs of preterm labor until E17.5. Beginning on E18.5, dams were housed singly, and cages were checked twice daily for evidence of pups. Gestational age at the time of delivery was recorded and was considered postnatal day 0. Twice daily, pups were monitored and the number of live and deceased pups were recorded until postnatal day 7. On postnatal days 4 and 7, pup weight was recorded. On postnatal day 7, pup intestines and liver were collected, homogenized, and plated on ChromAgar Orientation to quantify *E. coli* CFU. Dams were also euthanized on postnatal day 7, and maternal bladder, kidneys, vagina, and uterus were collected, homogenized, and plated as described above to quantify *E. coli* CFU.

### Immune cell profiling of maternal bladder, decidua, and spleen and fetal placenta

Maternal bladder and placenta were transected so that one-third tissue was homogenized as above for quantifying CFU, and two-thirds was processed for flow cytometric analysis. Maternal decidua from the two most proximal placenta were processed for flow cytometry, while the remaining decidua were homogenized. Bladder, decidua, and placenta were placed in RPMI and finely minced with scissors until the tissue could pass through a P1000 pipette tip. Collagenase (1 mg/mL, Sigma C5138-1G) and DNase (50 U/mL, Thermo Fisher J62229.MC) were added, then the samples were incubated at 37°C with shaking at 250 RPM for 30 minutes. 400 μL was then filtered through a 40 µm filter into 800 μL RPMI with 10% heat-inactivated FBS and stored on ice. The remaining tissue then went through a second digestion with fresh RPMI, collagenase, and DNase. From the second digestion, 400 μL was then filtered as above. Samples were then centrifuged at 500 × g for 10 minutes. Bladder tissue was resuspended in 200 μL, all of which was analyzed. Deciduae and placentae were resuspended in 500 μL, 50 μL of which was analyzed for each panel. Spleens were mechanically dissociated and filtered through a 40 µm filter into 2 mL RPMI, then centrifuged at 500 × g for 10 minutes. Spleens were then resuspended in 1 mL Red Cell Lysis Buffer (Biosearch Technologies MRC0912H-1) and incubated at room temperature for 10 minutes. After lysis, 9 mL sterile PBS was added and spleens were centrifuged. Pelleted spleens were resuspended in 5 mL sterile PBS, 50 μL of which was analyzed for each panel. Samples were then stained with 50 μL Zombie Aqua (1:1000 dilution) for 15 minutes at room temperature in the dark. Samples were then washed with PBS and pelleted via centrifugation. Samples were then resuspended in 50 μL Fc block (1:200 dilution) and incubated at 4°C in the dark for 15 minutes, followed by a PBS wash and centrifugation. Antibody cocktail panel 1 included: anti-CD45 (BV605, clone 30-F11, BD 563053), anti-CD11b (APC-Cy7 Clone M1/70 BD 561039), anti-CD11c (BV786, clone HL3 BD 563735), anti-Ly6G (AF700, clone 1A8, BD 561236), anti-NK1.1 (BUV737, clone PK136, BD 741715), anti-CD19 (BB515, clone 1D3, BD 564509), anti-CD3 (BV480, clone 145-2C11, BD 746368), anti-CD8 (PE-Cy7, clone QA17A07, Biolegend 155018), anti-CD4 (PE-CF594, clone RM4-5, BD 562314), anti-CD25 (APC, clone PC61, BD 557192), anti-FCεR1α (PE, clone MAR-1, BD 566995), and anti-CD64 (BV421, clone X54-5/7.1, Biolegend 139309). Antibody cocktail panel 2 included: anti-CD45 (BV605, clone 30-F11, BD 563053), anti-CD3 (BV480, clone 145-2C11, BD 746368), anti-CD8 (PE-Cy7, clone QA17A07, Biolegend 155018), anti-CD4 (PE-CF594, clone RM4-5, BD 562314). Intracellular panel included: anti-Tbet (BV711 clone 04-46 BD 563320), anti-GATA3 (AF647 clone l50-823, BD 560068), anti-FOXp3 (BV421 clone 3G3, BD 567458), anti-RORγT (BB515 clone Q31-378, BD 567175). Samples were then resuspended in 100 μL antibody cocktail Panel 1 or Panel 2and incubated at 4°C overnight for at least 16 hours. The following morning, samples stained with Panel 1 were washed with sterile PBS and resuspended in 200 μL FACS buffer (1% BSA, 0.5% EDTA, 0.05% sodium azide in PBS). Samples stained with Panel 2 were resuspended in 100 μL fixation buffer (Thermo Fisher GAS003) for 15 minutes at room temperature. Samples were then washed with PBS, centrifuged, and resuspended in the intracellular panel. After incubating for 30 minutes at room temperature in the dark, samples were washed, centrifuged, and resuspended in 200 μL FACS buffer for analysis. Data was acquired using a CytekBio Aurora and post-acquisition analyses were done using FlowJo software version 10.10.0. Gating strategy is shown in **Fig. S2**. Immune cell subsets were delineated from the CD45+ Zombie Aqua-population and defined as: dendritic cells (CD11c+CD11b-), neutrophils (CD11c-CD11b+Ly6G+), natural killer cells (CD11c-CD11b+Ly6G-CD64-NK1.1+), macrophages (CD11c-CD11b+Ly6G-CD64+NK1.1-), mast cells (CD11c-CD11b+Ly6G-CD64-NK1.1-FcεR1α+), B cells (CD11c-CD11b-CD19+CD3-), T cells (CD11c-CD11b-CD19-CD3+), CD4+ T cells (CD11c-CD11b-CD19-CD3+CD4+CD8-), CD8 positive T cells (CD11c-CD11b-CD19-CD3+CD4-CD8+), NKT cells (CD11c-CD11b-CD19-CD3+CD4-CD8-NK1.1+), FOXp3+ CD4+ T cells (CD3+CD4+CD8-FOXp3+), RORγT+ CD4+ T cells (CD3+CD4+CD8-RORγT+), T_H1_ T cells (CD3+CD4+CD8-RORγT-FOXp3-TBET+GATA3-), T_H2_ T cells (CD3+CD4+CD8-RORγT-FOXp3-TBET-GATA3+), T_H17_ T cells (CD3+CD4+CD8-RORγT+FOXp3-TBET-GATA3-), regulatory T cells (CD3+CD4+CD8-RORγT-FOXp3+TBET-GATA3-), FOXp3+RORγT+ T Cells (CD3+CD4+CD8-RORγT+FOXp3+TBET-GATA3-), and T_H0_ T cells (CD3+CD4+CD8-RORγT-FOXp3-TBET-GATA3-).

### Bulk RNA sequencing and analysis

Four hours post infection, maternal bladder and decidua and fetal placenta were dissected and placed in tubes containing 1.0mm zirconia/silica beads and 500 μL RNA protect (Qiagen). Tissues were homogenized using a MagNA Lyser instrument (Roche Diagnostics) for 60 seconds at 6,000 RPM. Homogenization was repeated for a total of three cycles. RNA extractions were performed on 10 mg (bladder) or 20 mg (decidua, placenta) by RNeasy Plus Mini Kit (Qiagen 74134). RNA sequencing was then performed by Novogene, where library construction and sequencing were performed via Illumina NovaSeq platform. Read mapping was performed using Hisat2 (v2.0.5) to reference genome GRCm39 and quantified using featureCounts (v1.5.0-p3). Raw count normalization and differential gene expression analysis was performed using R package DESeq2 (v1.42.0). R Studio (v2023.12.0-369) was used to generate heatmaps, PCA plots, volcano plots, and pathway enrichment plots. The R packages enhanced volcano and ashr LFC shrinkage were used for data visualization. fGSEA (v1.28.0) was used for gene set enrichment analysis with 10,000 permutations, a minimum gene set of 15 and maximum of 500 genes, and the Reactome gene set collection from the Molecular Signatures Database.

### Cytokine quantification in murine tissues

Bladder, decidual, and placental homogenates as well as maternal serum were analyzed for cytokine expression (IL-1α, IL-1β, IL-2, IL-3, IL-4, IL-5, IL-6, IL-9, IL-10, IL-12(p40), IL-12(p70), IL-13, IL-17A, Eotaxin, G-CSF, GM-CSF, IFN-γ, KC, MCP-1, MIP-1α, MIP-1β, RANTES, and TNF) via a 23-plex assay (Bio-Rad, M60009RDP). Samples were thawed on ice, then centrifuged at 10,000 × g for 10 minutes. Serum samples were diluted 1:4 in PBS. Samples were then processed according to manufacturer’s recommendations and analyzed on a Luminex Magpix with data analysis using Milliplex Analyst.

### Fetal sex determination

Fetal sex was determined via PCR targeting Rbm31x/y (F: 5’-CAC CTT AAG AAC AAG CCA ATA CA-3’, R: 5’-GGC TTG TCC TGA AAA CAT TTG G-3’) as described previously(86). 10 μL of placental lysate was boiled at 95°C for 10 minutes. 2 µL boiled lysate was then combined with 1 µL each of forward and reverse primers, 6 µL molecular grade water, and 10 µL 2X Platinum Hot Start PCR mastermix and PCR amplification was performed using the following conditions: 94°C for 2 min, followed by 30 cycles of 94°C for 20 s, 50°C for 20 s, 68°C for 30 s, and then 68°C for 5 min followed by a hold at 4°C. Amplification products were then run on a 1.3% agarose gel containing SYBR Safe stain (VWR 470193-138) and imaged on a UV transilluminator.

### Tissue fixation, staining, and histological scoring

After euthanasia, bladders were inflated with 10% neutral buffered formalin (VWR) via transurethral catheterization. Bladders were then tied off using embroidery thread, the catheter was removed, and the bladder was dissected out and placed in a histology cassette. The first placenta with intact decidua was dissected from each horn and placed in the histology cassette with the bladder for each dam. The cassettes were then fixed overnight in 10% neutral buffered formalin. After fixation, cassettes were transferred into 70% ethanol for storage until processing and embedding in paraffin. 3mm sections were cut 1/3 of the way into each paraffin block. Slides were stained with hematoxylin and eosin (H&E), assessed for inflammation (hemorrhage, edema, epithelial disruption, luminal or submucosal immune cell infiltration, apoptosis), and scored on a scale of 0 to 4 (no, mild, moderate, or severe inflammation) by a blinded veterinary pathologist.

### Recombinant IL-10 treatment

On E13.5, immediately after infection, dams were treated with 5 μg recombinant murine IL-10 (R&D Systems 417-ML-025/CF) resuspended in sterile PBS. IL-10 was administered as a 50 μL intraperitoneal injection, and 50 μL sterile PBS was used for controls. Mice were then monitored twice daily as described above, and euthanized when signs of preterm labor were detected or on E17.5. Maternal bladder, kidneys, vagina, and decidua and fetal placenta were homogenized and plated for CFU quantification as described above. Fetal weight was recorded.

### Cytokine quantification in human urine

Human subjects research, exempt from informed consent, was approved under University of South Alabama (USA) IRB, #2178590-1, and BCM IRB, protocol H-54361. Discarded urine samples from pregnant patients and nonpatient controls were obtained from the USA Hospital Clinical Microbiology Lab and stored at -20°C. Patient information, including birth outcomes, were collected retrospectively via chart review. Specimens were thawed overnight at 4°C, then centrifuged at 10,000 × g for 10 minutes. Samples were then diluted 1:2 in PBS and processed according to manufacturer recommendations. Samples were analyzed on a Luminex Magpix with data analysis using Milliplex Analyst. Samples were evaluated using the MILLIPLEX Human Cytokine/Chemokine/Growth Factor Panel A Magnetic Bead Panel (Millipore HCYTA-60K-PX48). Creatinine concentration was measured using the Creatinine Parameter colorimetric assay (R&D Systems KGE005). Samples were also evaluated for S100A9 concentration by ELISA (R&D Systems DY5578). All cytokines, chemokines, and growth factors were normalized to creatinine. sCD40L, Fractalkine, IFNg, IL-2, IL-7, IL-12 (p40), IL-17A, TNFb, IL-3, IL-13, IL-17E/IL-25, FGF-2, PDGF-AB/BB, MIP-1a, and MIP-1b were excluded from later analyses due to more than half of samples falling below the limit of detection. To develop predictive models, multiple logistic regression was performed in GraphPad Prism using a binary outcome of preterm birth (birth prior to 37 weeks gestation) or term birth (birth at 37 weeks or later gestation). Fitness of model was evaluated by area under the ROC curve.

### Statistics

All experiments were completed in at least duplicate unless otherwise indicated with results combined prior to analyses. Experimental sample size (n) and exact tests used are indicated in figure legends. Bacterial burdens and cytokine data were assessed by Mann-Whitney or Kruskal-Wallis tests when comparing two or three groups, respectively. Two-Way ANOVA with Benjamini, Krieger and Yekutieli false discovery rate correction for multiple comparisons (q<0.05) was employed for flow cytometric data and bacterial burdens for multiple tissues. Survival was analyzed using the log-rank Mantel-Cox test. Differential gene expression was determined using a generalized linear model where a Log_2_-fold change>1.5 and an adjusted p-value <0.05 were considered differentially expressed. Multivariable logistic regression analyses were evaluated using the area under the ROC curve with a p-value<0.05. Statistical analyses were performed using GraphPad Prism v10.2.3.

## Acknowledgements

We are grateful to the vivarium staff at BCM for animal husbandry, and Dr. Brian Simons for histopathological assessment of murine tissues. Flow cytometry was performed by the BCM Cytometry and Cell Sorting Core with funding from CPRIT award (CPRIT-RP180672) and NIH awards (CA125123, RR024574) with assistance from Claude Chew, Padmini Narayanan, and Joel Sederstrom. Multiplex cytokine analysis was performed by the BCM Antibody-based Proteomics Core, supported in part by CPRIT Proteomics & Metabolomics Core Facility Support Award (RP210227) and NCI Cancer Center Support Grant (P30CA125123). We thank Shixia Huang, Michael Nguyen, Zhongcheng Shi, and Yuan Yao for their excellent technical assistant in performing the Luminex experiments, data preliminary analyses and QC, and project consultation. We are grateful to the patients who contributed samples to this work, and the clinicians and scientists who contributed to sample collection and processing. SO, VME, MEM, and JJZ were supported by NIH F31 awards (HD111236, AI167547, AI167538, DK136201) respectively. Studies were supported by University of South Alabama College of Medicine start-up funds to AES and Burroughs Wellcome Next Gen Pregnancy and NIH R01 (DK128053) awards to KAP. The funders had no role in study design, data collection/interpretation, or the decision to publish.

## Author contributions

Research studies were designed by SO and KAP. SO, AL, VME, HB, JJZ, CS, MEM, ZH, RW, RCF, AES, and KAP conducted experiments and acquired data. Data were analyzed by SO and KAP. SO and KAP drafted the manuscript, and all authors contributed to manuscript review and edits.

## REFERENCES

1. H. Blencowe, S. Cousens, D. Chou, M. Oestergaard, L. Say, A. B. Moller, M. Kinney, J. Lawn, Born Too Soon: The global epidemiology of 15 million preterm births. Reprod Health 10, 2 (2013).

2. R. L. Goldenberg, J. F. Culhane, J. D. Iams, R. Romero, Epidemiology and causes of preterm birth. Lancet 371, 75–84 (2008).

3. R. J. Baer, N. Nidey, G. Bandoli, B. D. Chambers, C. D. Chambers, S. Feuer, D. Karasek, S. P. Oltman, L. Rand, K. K. Ryckman, L. L. Jelliffe-Pawlowski, Risk of Early Birth among Women with a Urinary Tract Infection: A Retrospective Cohort Study. AJP Rep 11, E5–E14 (2021).

4. E. Mazor-Dray, A. Levy, F. Schlaeffer, E. Sheiner, Maternal urinary tract infection: is it independently associated with adverse pregnancy outcome? Journal of Maternal-Fetal & Neonatal Medicine 22, 124–128 (2009).

5. N. Abou Heidar, J. Degheili, A. Yacoubian, R. Khauli, Management of urinary tract infection in women: A practical approach for everyday practice. Urol Ann 11, 339–346 (2019).

6. Y. Ansaldi, B. Martinez de Tejada Weber, Urinary tract infections in pregnancy. Clin Microbiol Infect 29, 1249–1253 (2023).

7. B. Abu-Raya, C. Michalski, M. Sadarangani, P. M. Lavoie, Maternal Immunological Adaptation During Normal Pregnancy. Front Immunol 11 (2020), doi:10.3389/FIMMU.2020.575197.

8. N. Aghaeepour, E. A. Ganio, D. Mcilwain, A. S. Tsai, M. Tingle, S. Van Gassen, D. K. Gaudilliere, Q. Baca, L. McNeil, R. Okada, M. S. Ghaemi, D. Furman, R. J. Wong, V. D. Winn, M. L. Druzin, Y. Y. El-Sayed, C. Quaintance, R. Gibbs, G. L. Darmstadt, G. M. Shaw, D. K. Stevenson, R. Tibshirani, G. P. Nolan, D. B. Lewis, M. S. Angst, B. Gaudilliere, An immune clock of human pregnancy. Sci Immunol 2, undefined-undefined (2017).

9. G. Mor, P. Aldo, A. B. Alvero, The unique immunological and microbial aspects of pregnancy. Nature Reviews Immunology 2017 17:8 17, 469–482 (2017).

10. Jr. John E. Delzell, M. L. Lefevre, Urinary Tract Infections During Pregnancy. Am Fam Physician 61, 713–720 (2000).

11. R. Cohen, G. Gutvirtz, T. Wainstock, E. Sheiner, Maternal urinary tract infection during pregnancy and long-term infectious morbidity of the offspring. Early Hum Dev 136, 54–59 (2019).

12. A. K. Kaul, S. Khan, M. G. Martens, J. T. Crosson, V. R. Lupo, R. Kaul, Experimental gestational pyelonephritis induces preterm births and low birth weights in C3H/HeJ mice. Infect Immun 67, 5958–5966 (1999).

13. M. Bolton, D. J. Horvath, B. Li, H. Cortado, D. Newsom, P. White, S. Partida-Sanchez, S. S. Justice, Intrauterine growth restriction is a direct consequence of localized maternal uropathogenic Escherichia coli cystitis. PLoS One 7 (2012), doi:10.1371/JOURNAL.PONE.0033897.

14. S. Henry, S. M. Lewis, S. L. Cyrill, M. K. Callaway, D. Chatterjee, A. V. Hanasoge Somasundara, G. Jones, X. Y. He, G. Caligiuri, M. F. Ciccone, I. A. Diaz, A. A. Biswas, E. Hernandez, T. Ha, J. E. Wilkinson, M. Egeblad, D. A. Tuveson, C. O. dos Santos, Host response during unresolved urinary tract infection alters female mammary tissue homeostasis through collagen deposition and TIMP1. Nat Commun 15 (2024), doi:10.1038/S41467-024-47462-7.

15. N. Gomez-Lopez, D. StLouis, M. A. Lehr, E. N. Sanchez-Rodriguez, M. Arenas-Hernandez, Immune cells in term and preterm labor. Cell Mol Immunol 11, 571–581 (2014).

16. M. J. Allard, A. Giraud, M. Segura, G. Sebire, Sex-specific maternofetal innate immune responses triggered by group B Streptococci. Sci Rep 9 (2019), doi:10.1038/S41598-019-45029-X.

17. Q. Na, A. Chudnovets, J. Liu, J. Y. Lee, J. Dong, N. Shin, N. Elsayed, J. Lei, I. Burd, Placental Macrophages Demonstrate Sex-Specific Response to Intrauterine Inflammation and May Serve as a Marker of Perinatal Neuroinflammation. J Reprod Immunol 147 (2021), doi:10.1016/j.jri.2021.103360.

18. L. L. Shook, K. E. James, D. J. Roberts, C. E. Powe, R. H. Perlis, K. L. Thornburg, P. F. O’Tierney-Ginn, A. G. Edlow, Sex-specific impact of maternal obesity on fetal placental macrophages and cord blood triglycerides. Placenta 140, 100–108 (2023).

19. A. N. Battarbee, A. V. Glover, C. J. Vladutiu, C. Gyamfi-Bannerman, S. Aliaga, T. A. Manuck, K. A. Boggess, Sex-Specific Differences in Late Preterm Neonatal Outcomes. Am J Perinatol 36, 1223–1228 (2019).

20. S. Galjaard, L. Ameye, C. C. Lees, A. Pexsters, T. Bourne, D. Timmerman, R. Devlieger, Sex differences in fetal growth and immediate birth outcomes in a low-risk Caucasian population. Biol Sex Differ 10, 48 (2019).

21. M. A. Mulvey, J. D. Schilling, S. J. Hultgren, Establishment of a persistent Escherichia coli reservoir during the acute phase of a bladder infection. Infect Immun 69, 4572–4579 (2001).

22. D. Yu, G. Banting, N. F. Neumann, A review of the taxonomy, genetics, and biology of the genus Escherichia and the type species Escherichia coli. Can J Microbiol 67, 553–571 (2021).

23. E. Sheiner, E. Mazor-Drey, A. Levy, Asymptomatic bacteriuria during pregnancy. J Matern Fetal Neonatal Med 22, 423–427 (2009).

24. V. Mercado-Evans, C. Chew, C. Serchejian, A. Saltzman, M. E. Mejia, J. J. Zulk, I. Cornax, V. Nizet, K. A. Patras, Tamm-Horsfall protein augments neutrophil NETosis during urinary tract infection. bioRxiv (2024), doi:10.1101/2024.02.01.578501.

25. V. D. Radu, R. C. Costache, P. Onofrei, L. Antohi, R. L. Bobeica, I. Linga, I. Tanase-Vasilache, A. I. Ristescu, A. M. Murgu, I. L. Miftode, B. A. Stoica, Factors Associated with Increased Risk of Urosepsis during Pregnancy and Treatment Outcomes, in a Urology Clinic. Medicina (Kaunas) 59 (2023), doi:10.3390/MEDICINA59111972.

26. I. Abu Aleinein, E. Salem Sokhn, Knowledge and prevalence of urinary tract infection among pregnant women in Lebanon. Heliyon 10 (2024), doi:10.1016/J.HELIYON.2024.E37277.

27. B. Lengelé, P. Scalliet, Anatomical bases for the radiological delineation of lymph node areas. Part III: Pelvis and lower limbs. Radiotherapy and Oncology 92, 22–33 (2009).

28. F. Wynne, M. Ball, A. S. McLellan, P. Dockery, W. Zimmermann, T. Moore, Mouse pregnancy-specific glycoproteins: tissue-specific expression and evidence of association with maternal vasculature. Reproduction 131, 721–732 (2006).

29. J. Kawai, A. Shinagawa, K. Shibata, M. Yoshino, M. Itoh, Y. Ishii, T. Arakawa, A. Hara, Y. Fukunishi, H. Konno, J. Adachi, S. Fukuda, K. Aizawa, M. Izawa, K. Nishi, H. Kiyosawa, S. Kondo, I. Yamanaka, T. Saito, Y. Okazaki, T. Gojobori, H. Bono, T. Kasukawa, R. Saito, K. Kadota, H. Matsuda, M. Ashburner, S. Batalov, T. Casavant, W. Fleischmann, T. Gaasterland, C. Gissi, B. King, H. Kochiwa, P. Kuehl, S. Lewis, Y. Matsuo, I. Nikaido, G. Pesole, J. Quackenbush, L. M. Schriml, F. Staubli, R. Suzuki, M. Tomita, L. Wagner, T. Washio, K. Sakai, T. Okido, M. Furuno, H. Aono, R. Baldarelli, G. Barsh, J. Blake, D. Boffelli, N. Bojunga, P. Carninci, M. F. De Bonaldo, M. J. Brownstein, C. Bult, C. Fletcher, M. Fujita, M. Gariboldi, S. Gustincich, D. Hill, M. Hofmann, D. A. Hume, M. Kamiya, N. H. Lee, P. Lyons, L. Marchionni, J. Mashima, J. Mazzarelli, P. Mombaerts, P. Nordone, B. Ring, M. Ringwald, I. Rodriguez, N. Sakamoto, H. Sasaki, K. Sato, C. Schönbach, T. Seya, Y. Shibata, K. F. Storch, H. Suzuki, K. Toyo-Oka, K. H. Wang, C. Weitz, C. Whittaker, L. Wilming, A. Wynshaw-Boris, K. Yoshida, Y. Hasegawa, H. Kawaji, S. Kohtsuki, Y. Hayashizaki, Functional annotation of a full-length mouse cDNA collection. Nature 409, 685–689 (2001).

30. J. Wessells, D. Wessner, R. Parsells, K. White, D. Finkenzeller, W. Zimmermann, G. Dveksler, Pregnancy specific glycoprotein 18 induces IL-10 expression in murine macrophages., doi:10.1002/1521-4141(200007)30:7.

31. R. Kessous, A. Y. Weintraub, R. Sergienko, T. Lazer, F. Press, A. Wiznitzer, E. Sheiner, Bacteruria with group-B *streptococcus*: is it a risk factor for adverse pregnancy outcomes? J Matern Fetal Neonatal Med 25, 1983–1986 (2012).

32. W. A. Bayih, M. Y. Ayalew, E. S. Chanie, B. B. Abate, S. A. Alemayehu, D. M. Belay, Y. A. Aynalem, D. A. Sewyew, S. D. Kebede, A. Demis, G. Y. Yitbarek, M. A. Tassew, B. M. Birhan, A. Y. Alemu, The burden of neonatal sepsis and its association with antenatal urinary tract infection and intra-partum fever among admitted neonates in Ethiopia: A systematic review and meta-analysis. Heliyon 7 (2021), doi:10.1016/J.HELIYON.2021.E06121.

33. M. J. Patrick, Influence of maternal renal infection on the foetus and infant. Arch Dis Child 42, 208–213 (1967).

34. M. A. Rafi, M. M. Z. Miah, M. A. Wadood, M. G. Hossain, Risk factors and etiology of neonatal sepsis after hospital delivery: A case-control study in a tertiary care hospital of Rajshahi, Bangladesh. PLoS One 15 (2020), doi:10.1371/JOURNAL.PONE.0242275.

35. M. Liu, X. Luo, Q. Xu, H. Yu, L. Gao, R. Zhou, T. Wang, Adipsin of the Alternative Complement Pathway Is a Potential Predictor for Preeclampsia in Early Pregnancy. Front Immunol 12 (2021), doi:10.3389/FIMMU.2021.702385.

36. Z. A. Broere-Brown, M. C. Adank, L. Benschop, M. Tielemans, T. Muka, R. Gonçalves, W. M. Bramer, J. D. Schoufour, T. Voortman, E. A. P. Steegers, O. H. Franco, S. Schalekamp-Timmermans, Fetal sex and maternal pregnancy outcomes: a systematic review and meta-analysis. Biol Sex Differ 11 (2020), doi:10.1186/S13293-020-00299-3.

37. I. Ambite, D. Butler, M. L. Y. Wan, T. Rosenblad, T. H. Tran, S. M. Chao, C. Svanborg, Molecular determinants of disease severity in urinary tract infection. Nat Rev Urol 18, 468–486 (2021).

38. C. Petersson, S. Hedges, K. Stenqvist, T. Sandberg, H. Connell, C. Svanborg, Suppressed antibody and interleukin-6 responses to acute pyelonephritis in pregnancy. Kidney Int 45, 571–577 (1994).

39. M. Azami, Z. Jaafari, M. Masoumi, M. Shohani, G. Badfar, L. Mahmudi, S. Abbasalizadeh, The etiology and prevalence of urinary tract infection and asymptomatic bacteriuria in pregnant women in Iran: a systematic review and Meta-analysis. BMC Urol 19 (2019), doi:10.1186/S12894-019-0454-8.

40. L. D. Mera-Lojano, L. A. Mejía-Contreras, S. M. Cajas-Velásquez, S. J. Guarderas-Muñoz, [Prevalence and risk factors of urinary tract infection in pregnant women]. Rev Med Inst Mex Seguro Soc 61, 590–596 (2023).

41. E. Zhang, Q. Zeng, Y. Xu, J. Lu, C. Li, K. Xiao, X. Li, J. Li, T. Li, C. Li, L. Zhang, A smartphone-based immunochromatographic strip platform for on-site quantitative detection of antigenic targets. Lab Chip 24 (2024), doi:10.1039/D4LC00484A.

42. X. Zhao, A. Xu, X. Lu, B. Chen, Y. Hua, Y. Ma, Association of phthalates exposure and sex steroid hormones with late-onset preeclampsia: a case-control study. BMC Pregnancy Childbirth 24 (2024), doi:10.1186/S12884-024-06793-5.

43. L. S. Lentz, A. J. Stutz, N. Meyer, K. Schubert, I. Karkossa, M. von Bergen, A. C. Zenclussen, A. Schumacher, Human chorionic gonadotropin promotes murine Treg cells and restricts pregnancy-harmful proinflammatory Th17 responses. Front Immunol 13 (2022), doi:10.3389/FIMMU.2022.989247.

44. A. Sen, A. Kaul, R. Kaul, Estrogen receptors in human bladder cells regulate innate cytokine responses to differentially modulate uropathogenic *E. coli* colonization. Immunobiology 226 (2021), doi:10.1016/J.IMBIO.2020.152020.

45. S. I. Ramírez, E. A. Suniega, M. I. Laughrey, Endocrinology During Pregnancy. Prim Care 51, 535–547 (2024).

46. F. F. Martínez, C. P. Knubel, M. C. Sánchez, L. Cervi, C. C. Motrán, Pregnancy-specific glycoprotein 1a activates dendritic cells to provide signals for Th17-, Th2-, and Treg-cell polarization. Eur J Immunol 42, 1573–1584 (2012).

47. S. K. Snyder, D. H. Wessner, J. L. Wessells, R. M. Waterhouse, L. M. Wahl, W. Zimmermann, G. S. Dveksler, Pregnancy-specific glycoproteins function as immunomodulators by inducing secretion of IL-10, IL-6 and TGF-beta1 by human monocytes. Am J Reprod Immunol 45, 205–216 (2001).

48. D. K. Shanley, P. A. Kiely, K. Golla, S. Allen, K. Martin, R. T. O’Riordan, M. Ball, J. D. Aplin, B. B. Singer, N. Caplice, N. Moran, T. Moore, Pregnancy-specific glycoproteins bind integrin αIIbβ3 and inhibit the platelet-fibrinogen interaction. PLoS One 8 (2013), doi:10.1371/JOURNAL.PONE.0057491.

49. T. E. C. Kieffer, A. Laskewitz, S. A. Scherjon, M. M. Faas, J. R. Prins, Memory T Cells in Pregnancy. Front Immunol 10 (2019), doi:10.3389/FIMMU.2019.00625.

50. A. M. Mitchell, M. Palettas, L. M. Christian, Fetal sex is associated with maternal stimulated cytokine production, but not serum cytokine levels, in human pregnancy. Brain Behav Immun 60, 32–37 (2017).

51. A. H. Jarmund, G. F. Giskeødegård, M. Ryssdal, B. Steinkjer, L. M. T. Stokkeland, T. S. Madssen, S. N. Stafne, S. Stridsklev, T. Moholdt, R. Heimstad, E. Vanky, A. C. Iversen, Cytokine Patterns in Maternal Serum From First Trimester to Term and Beyond. Front Immunol 12 (2021), doi:10.3389/FIMMU.2021.752660.

52. A. Olmos-Ortiz, A. Olivares-Huerta, J. García-Quiroz, E. Avila, A. Halhali, B. Quesada-Reyna, F. Larrea, V. Zaga-Clavellina, L. Díaz, Cord Serum Calcitriol Inversely Correlates with Maternal Blood Pressure in Urinary Tract Infection-Affected Pregnancies: Sex-Dependent Immune Implications. Nutrients 13 (2021), doi:10.3390/NU13093114.

53. A. Olmos-Ortiz, A. Olivares-Huerta, J. García-Quiroz, T. Zariñán, R. Chavira, V. Zaga-Clavellina, E. Avila, A. Halhali, M. Durand, F. Larrea, L. Díaz, Placentas associated with female neonates from pregnancies complicated by urinary tract infections have higher cAMP content and cytokines expression than males. Am J Reprod Immunol 86 (2021), doi:10.1111/AJI.13434.

54. K. J. Baines, R. C. West, Sex differences in innate and adaptive immunity impact fetal, placental, and maternal health†. Biol Reprod 109, 256–270 (2023).

55. H. C. Osman, R. Moreno, D. Rose, M. E. Rowland, A. V. Ciernia, P. Ashwood, Impact of maternal immune activation and sex on placental and fetal brain cytokine and gene expression profiles in a preclinical model of neurodevelopmental disorders. J Neuroinflammation 21 (2024), doi:10.1186/S12974-024-03106-7.

56. P. Pantazi, M. Kaforou, Z. Tang, V. M. Abrahams, A. McArdle, S. Guller, B. Holder, Placental macrophage responses to viral and bacterial ligands and the influence of fetal sex. iScience 25, 105653 (2022).

57. S. A. Robertson, R. J. Skinner, A. S. Care, Essential role for IL-10 in resistance to lipopolysaccharide-induced preterm labor in mice. J Immunol 177, 4888–4896 (2006).

58. M. Busse, K. N. J. Campe, D. Nowak, A. Schumacher, S. Plenagl, S. Langwisch, G. Tiegs, A. Reinhold, A. C. Zenclussen, IL-10 producing B cells rescue mouse fetuses from inflammation-driven fetal death and are able to modulate T cell immune responses. Sci Rep 9 (2019), doi:10.1038/S41598-019-45860-2.

59. S. A. Robertson, A. S. Care, R. J. Skinner, Interleukin 10 regulates inflammatory cytokine synthesis to protect against lipopolysaccharide-induced abortion and fetal growth restriction in mice. Biol Reprod 76, 738–748 (2007).

60. J. C. Wommack, R. J. Ruiz, C. N. Marti, R. P. Stowe, C. E. L. Brown, C. Murphey, Interleukin-10 predicts preterm birth in acculturated Hispanics. Biol Res Nurs 15, 78–85 (2013).

61. R. J. Ruiz, N. Jallo, C. Murphey, C. N. Marti, E. Godbold, R. H. Pickler, Second trimester maternal plasma levels of cytokines IL-1Ra, Il-6 and IL-10 and preterm birth. J Perinatol 32, 483–490 (2012).

62. J. H. Rowe, J. M. Ertelt, M. N. Aguilera, M. A. Farrar, S. S. Way, Foxp3(+) regulatory T cell expansion required for sustaining pregnancy compromises host defense against prenatal bacterial pathogens. Cell Host Microbe 10, 54–64 (2011).

63. M. T. Rasquinha, M. Sur, N. Lasrado, J. Reddy, IL-10 as a Th2 Cytokine: Differences Between Mice and Humans. J Immunol 207, 2205–2215 (2021).

64. M. Impis Oglou, I. Tsakiridis, A. Mamopoulos, I. Kalogiannidis, A. Athanasiadis, T. Dagklis, Cervical length screening for predicting preterm birth: A comparative review of guidelines. J Clin Ultrasound 51, 472–478 (2023).

65. Z. A. Oskovi Kaplan, A. S. Ozgu-Erdinc, Prediction of Preterm Birth: Maternal Characteristics, Ultrasound Markers, and Biomarkers: An Updated Overview. J Pregnancy 2018 (2018), doi:10.1155/2018/8367571.

66. M. Nadeau-Vallée, D. Obari, C. Quiniou, W. D. Lubell, D. M. Olson, S. Girard, S. Chemtob, A critical role of interleukin-1 in preterm labor. Cytokine Growth Factor Rev 28, 37–51 (2016).

67. Y. Hatakeyama, H. Miura, A. Sato, Y. Onodera, N. Sato, D. Shimizu, Y. Kumazawa, H. Sanada, H. Hirano, Y. Terada, Neutrophil elastase in amniotic fluid as a predictor of preterm birth after emergent cervical cerclage. Acta Obstet Gynecol Scand 95, 1136–1142 (2016).

68. F. Namba, S. Ina, H. Kitajima, H. Yoshio, K. Mimura, S. Saito, I. Yanagihara, Annexin A2 in amniotic fluid: Correlation with histological chorioamnionitis, preterm premature rupture of membranes, and subsequent preterm delivery. Journal of Obstetrics and Gynaecology Research 38, 137–144 (2012).

69. Z. Alfirevic, K. Navaratnam, F. Mujezinovic, Amniocentesis and chorionic villus sampling for prenatal diagnosis. Cochrane Database Syst Rev 9 (2017), doi:10.1002/14651858.CD003252.PUB2.

70. P. Pruski, G. D. S. Correia, H. V. Lewis, K. Capuccini, P. Inglese, D. Chan, R. G. Brown, L. Kindinger, Y. S. Lee, A. Smith, J. Marchesi, J. A. K. McDonald, S. Cameron, K. Alexander-Hardiman, A. L. David, S. J. Stock, J. E. Norman, V. Terzidou, T. G. Teoh, L. Sykes, P. R. Bennett, Z. Takats, D. A. MacIntyre, Direct on-swab metabolic profiling of vaginal microbiome host interactions during pregnancy and preterm birth. Nat Commun 12 (2021), doi:10.1038/S41467-021-26215-W.

71. Z. Shaffer, R. Romero, A. L. Tarca, J. Galaz, M. Arenas-Hernandez, D. W. Gudicha, T. Chaiworapongsa, E. Jung, M. Suksai, K. R. Theis, N. Gomez-Lopez, The vaginal immunoproteome for the prediction of spontaneous preterm birth: A retrospective longitudinal study. Elife 13 (2024), doi:10.7554/ELIFE.90943.

72. K. Leitner, M. Al Shammary, M. Mclane, M. V. Johnston, M. A. Elovitz, I. Burd, IL-1 receptor blockade prevents fetal cortical brain injury but not preterm birth in a mouse model of inflammation-induced preterm birth and perinatal brain injury. Am J Reprod Immunol 71, 418–426 (2014).

73. T. A. Ayash, S. Y. Vancolen, M. Segura, M. J. Allard, G. Sebire, Protective Effects of Interleukin-1 Blockade on Group B *Streptococcus*-Induced Chorioamnionitis and Subsequent Neurobehavioral Impairments of the Offspring. Front Endocrinol (Lausanne) 13 (2022), doi:10.3389/FENDO.2022.833121.

74. Y. Takahashi, T. Takahashi, H. Usuda, S. Carter, E. L. Fee, L. Furfaro, S. Chemtob, D. M. Olson, J. A. Keelan, S. Kallapur, M. W. Kemp, Pharmacological blockade of the interleukin-1 receptor suppressed Escherichia coli lipopolysaccharide-induced neuroinflammation in preterm fetal sheep. Am J Obstet Gynecol MFM 5 (2023), doi:10.1016/J.AJOGMF.2023.101124.

75. T. Habelrih, D. É. Tremblay, E. Di Battista, X. Hou, A. Reuben, B. Ferri, S. E. Loiselle, F. Côté, P. Abram, W. D. Lubell, K. B. Leimert, C. Quiniou, S. Girard, D. M. Olson, S. Chemtob, Pharmacodynamic characterization of rytvela, a novel allosteric anti-inflammatory therapeutic, to prevent preterm birth and improve fetal and neonatal outcomes. Am J Obstet Gynecol 228, 467.e1–467.e16 (2023).

76. M. T. Aung, Y. Yu, K. K. Ferguson, D. E. Cantonwine, L. Zeng, T. F. McElrath, S. Pennathur, B. Mukherjee, J. D. Meeker, Prediction and associations of preterm birth and its subtypes with eicosanoid enzymatic pathways and inflammatory markers. Sci Rep 9 (2019), doi:10.1038/S41598-019-53448-Z.

77. B. M. Welch, A. P. Keil, J. P. Buckley, A. M. Calafat, K. E. Christenbury, S. M. Engel, K. M. O’Brien, E. M. Rosen, T. James-Todd, A. R. Zota, K. K. Ferguson, A. N. Alshawabkeh, J. F. Cordero, J. D. Meeker, E. S. Barrett, N. R. Bush, R. H. N. Nguyen, S. Sathyanarayana, S. H. Swan, D. E. Cantonwine, T. F. McElrath, J. Aalborg, D. Dabelea, A. P. Starling, R. Hauser, C. Messerlian, Y. Zhang, A. Bradman, B. Eskenazi, K. G. Harley, N. Holland, M. S. Bloom, R. B. Newman, A. G. Wenzel, J. M. Braun, B. P. Lanphear, K. Yolton, P. Factor-Litvak, J. B. Herbstman, V. A. Rauh, E. Z. Drobnis, A. E. Sparks, J. B. Redmon, C. Wang, A. M. Binder, K. B. Michels, D. D. Baird, A. M. Z. Jukic, C. R. Weinberg, A. J. Wilcox, D. Q. Rich, B. Weinberger, V. Padmanabhan, D. J. Watkins, I. Hertz-Picciotto, R. J. Schmidt, Associations Between Prenatal Urinary Biomarkers of Phthalate Exposure and Preterm Birth: A Pooled Study of 16 US Cohorts. JAMA Pediatr 176, 895–905 (2022).

78. S. M. Eick, S. D. Geiger, A. Alshawabkeh, M. Aung, E. S. Barrett, N. Bush, K. N. Carroll, J. F. Cordero, D. E. Goin, K. K. Ferguson, L. G. Kahn, D. Liang, J. D. Meeker, G. L. Milne, R. H. N. Nguyen, A. M. Padula, S. Sathyanarayana, K. R. Taibl, S. L. Schantz, T. J. Woodruff, R. Morello-Frosch, Urinary oxidative stress biomarkers are associated with preterm birth: an Environmental Influences on Child Health Outcomes program study. Am J Obstet Gynecol 228, 576.e1–576.e22 (2023).

79. Biomarkers for preterm births using non-invasive samples | NYU Langone Health (available at https://clinicaltrials.med.nyu.edu/clinicaltrial/1236/biomarkers-preterm-births-using/).

80. S. J. Fortunato, R. Menon, S. J. Lombardi, IL-15, a novel cytokine produced by human fetal membranes, is elevated in preterm labor. Am J Reprod Immunol 39, 16–23 (1998).

81. S. M. Gordon, Interleukin-15 in Outcomes of Pregnancy. Int J Mol Sci 22 (2021), doi:10.3390/IJMS222011094.

82. I. Mitrogiannis, E. Evangelou, A. Efthymiou, T. Kanavos, E. Birbas, G. Makrydimas, S. Papatheodorou, Risk factors for preterm birth: an umbrella review of meta-analyses of observational studies. BMC Medicine 2023 21:1 21, 1–17 (2023).

83. J. M. Bates, H. M. Raffi, K. Prasadan, R. Mascarenhas, Z. Laszik, N. Maeda, S. J. Hultgren, S. Kumar, Tamm-Horsfall protein knockout mice are more prone to urinary tract infection: rapid communication. Kidney Int 65, 791–797 (2004).

84. J. J. Zulk, J. R. Clark, S. Ottinger, M. B. Ballard, M. E. Mejia, V. Mercado-Evans, E. R. Heckmann, B. C. Sanchez, B. W. Trautner, A. W. Maresso, K. A. Patras, Phage Resistance Accompanies Reduced Fitness of Uropathogenic Escherichia coli in the Urinary Environment. mSphere 7 (2022), doi:10.1128/MSPHERE.00345-22/ASSET/074C0EF2-7EC1-44A2-8FBB-1A3C646354FB/ASSETS/IMAGES/MEDIUM/MSPHERE.00345-22-F007.GIF.

85. K. A. Patras, A. Coady, P. Babu, S. R. Shing, A. D. Ha, E. Rooholfada, S. L. Brandt, M. Geriak, R. L. Gallo, V. Nizet, Host Cathelicidin Exacerbates Group B *Streptococcus* Urinary Tract Infection. mSphere 5 (2020), doi:10.1128/MSPHERE.00932-19.

86. S. J. Tunster, Genetic sex determination of mice by simplex PCR. Biol Sex Differ 8, 6–9 (2017).

