## Supplemental Material for "Distinct maternofetal immune signatures delineate preterm birth onset following urinary tract infection"

**Contents:**

Supplementary Figures 1-5

Supplementary Tables 2-3

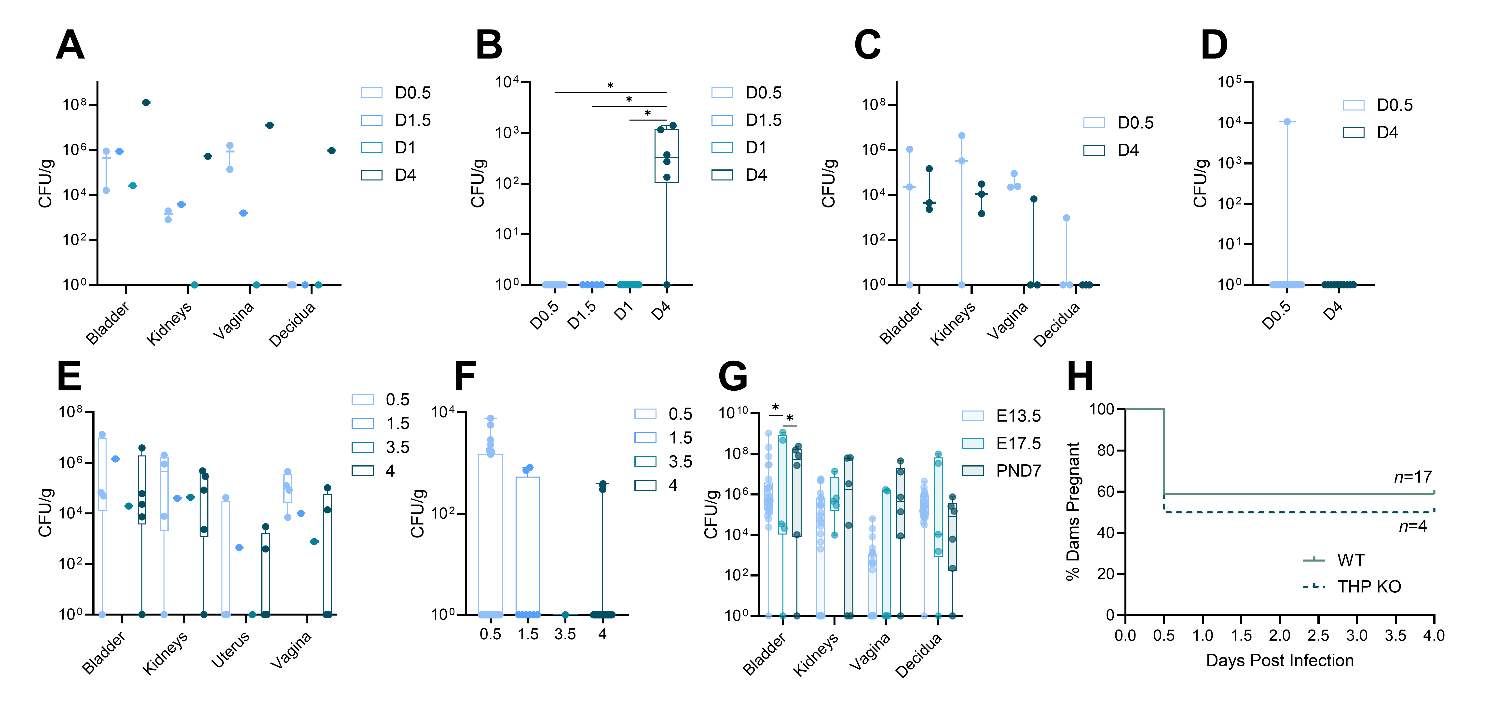

**Supplemental Figure 1. Bacterial burdens for SL2, SL323, GBS, and postnatal day 7 dams, and the impact of THP on preterm birth incidence.** Bladder, kidney, vaginal, and decidual (**A**) and placental (**B**) bacterial burdens for dams infected with SL2. Bladder, kidney, vaginal, and decidual (**C**) and placental (**D**) bacterial burdens for dams infected with SL323. Bladder, kidney, vaginal, and decidual (**E**) and placental (**F**) bacterial burdens for dams infected with GBS. Bladder, kidney, vaginal, and decidual bacterial burdens for dams infected with UTI89 on E13.5, E17.5, and postnatal day 7 (**G**). Kaplan-Meyer curve depicting percent of dams still pregnant in days post-infection for wildtype and Tamm-Horsfall Protein knockout (THP KO) mice (**H**). Experiments were performed at least twice with data combined. E13.5 and E17.5 data subsets (G) are also shown in Fig. 3A, 4A, and 6E. *n*=1-2 (A), *n*=5-15 (B), *n*=3 (C), *n*=10-13 (D), *n*=1-5 (E), *n*=1-36 (F), *n*=6 (G), *n*=4-17 (H). Box and whisker plots show median, all points, and extend from 25^th^ to 75^th^ percentiles. Data were analyzed by Mantel-Cox test (H), Kruskal-Wallis test (B,D,F), or Two-Way ANOVA with Benjamini, Krieger and Yekutieli correction for false discovery with a false discovery rate set at 5% (G). Some panels were not analyzed due to small sample size (A,C,E). *p<0.05.

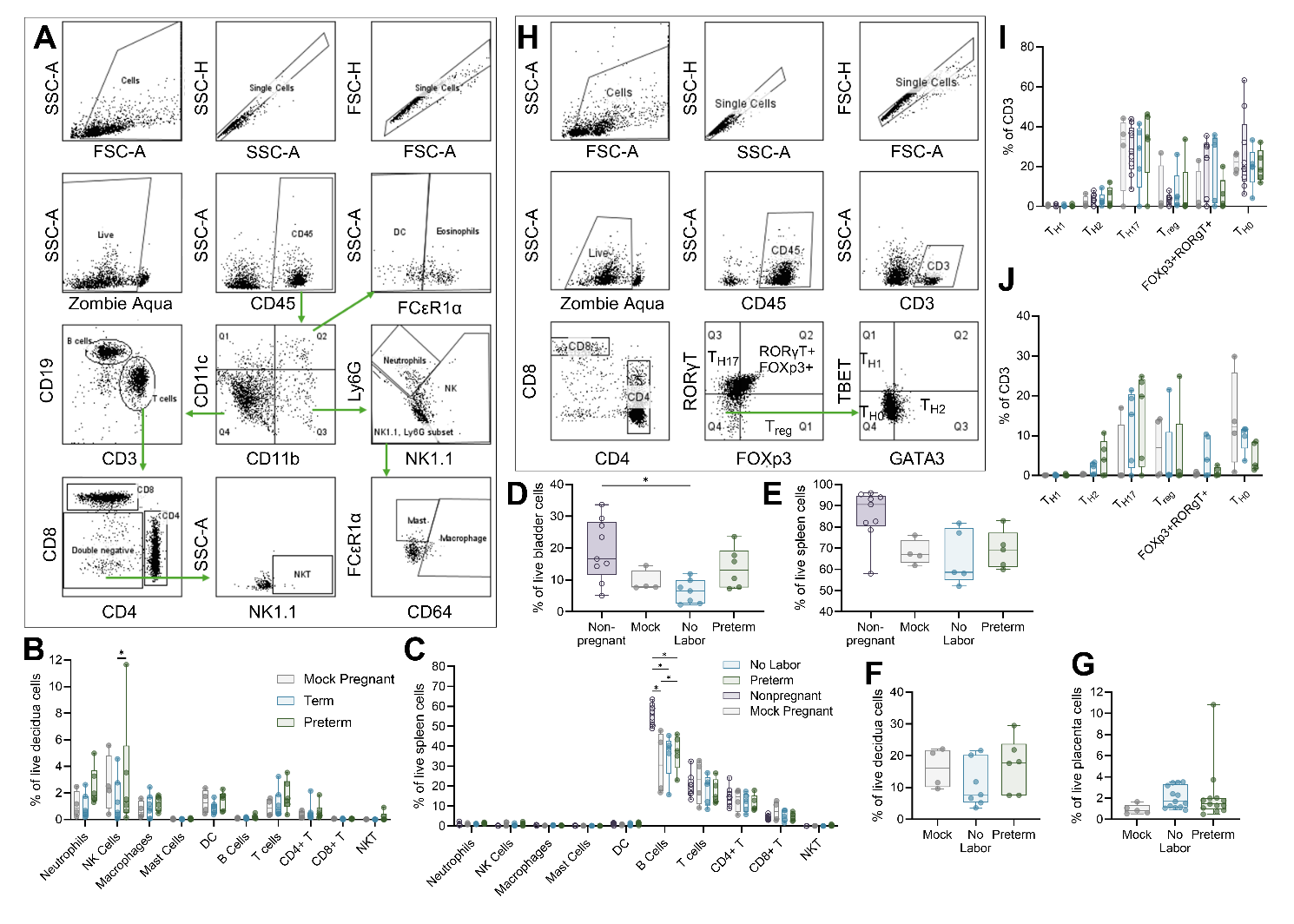

**Supplemental Figure 2. Flow cytometry gating schematics and additional immune cell profiling data.** Gating strategy for assessing immune cell composition (**A**). Immune cell phenotyping in the decidua (**B**) and spleen (**C**). CD45+ proportion of single cells in the bladder (**D**), spleen (**E**), decidua (**F**), and placenta (**G**). Gating strategy for assessing T helper lymphocyte phenotype (**H**). T helper phenotyping results for spleen (**I**) and decidua (**J**). Experiments were performed at least twice with data combined. *n*=4-7 (B), *n*=4-8 (C), *n*=4-9 (D,E,I), *n*=4-7 (F), *n*=5-13 (G), *n*=4-5 (J). Box and whisker plots show median, all points, and extend from 25^th^ to 75^th^ percentiles. Data were analyzed by Kruskal-Wallis test (D,E,F,G) or Two-Way ANOVA with Benjamini, Krieger and Yekutieli correction for false discovery with a false discovery rate set at 5% (B,C,I,J). *q<0.05.

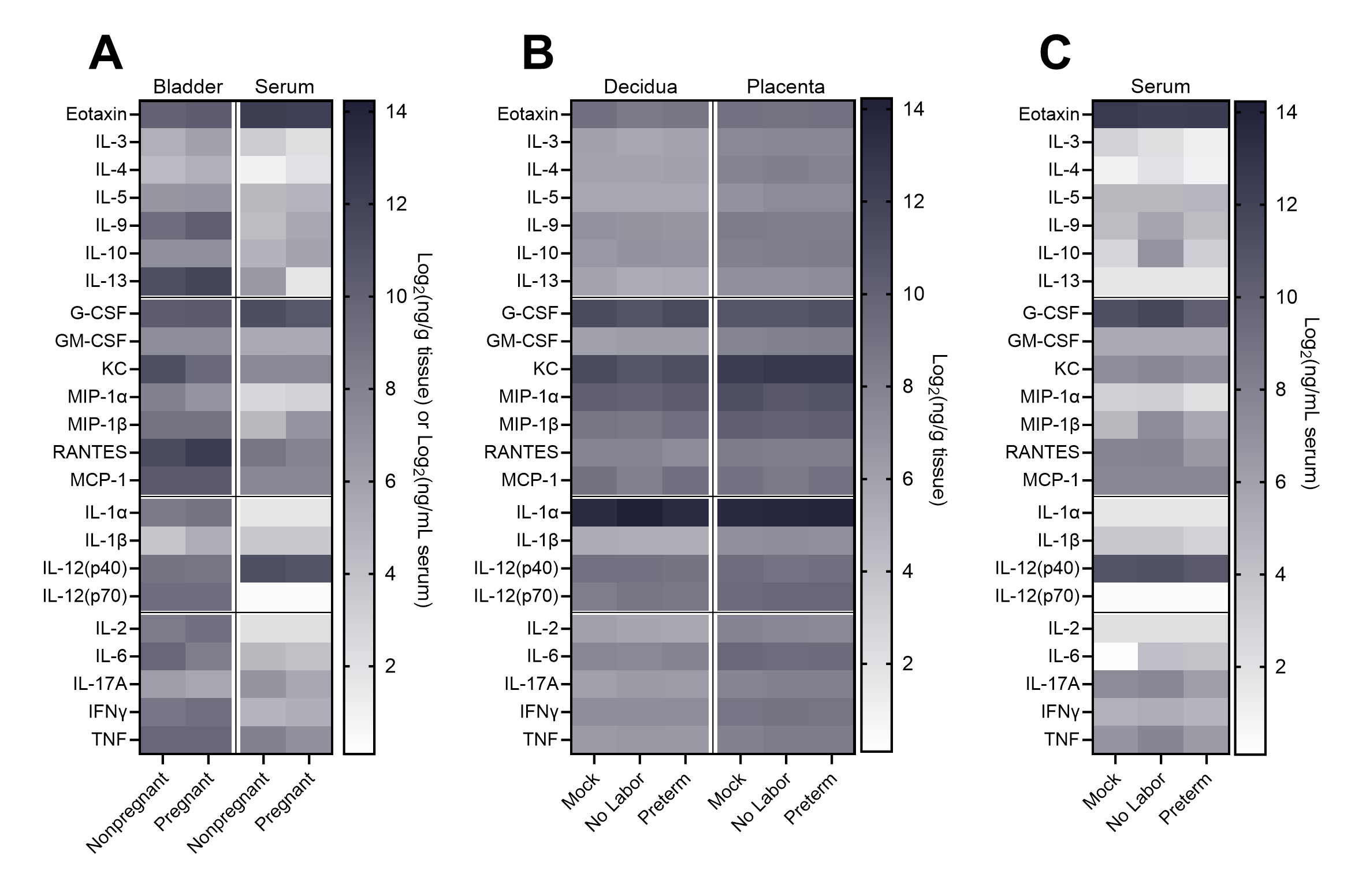
**Supplementary Figure 3. Heatmaps of cytokine quantitation in murine tissues.** Heatmap of 23 cytokines measured in the bladder and serum of nonpregnant and pregnant infected mice four hours post-infection (**A**). Heatmap of 23 cytokines measured in the decidua and placenta of mock and infected pregnant mice four hours post-infection (**B**). Heatmap of 23 cytokines measured in the serum of mock and infected pregnant mice four hours post-infection (**C**). Heatmaps show median value for each group. *n*=13-17 (A), *n*=7-19 (B), *n*=7-14 (C). Supplemental data to Figures 3, 4, 5, and 6.

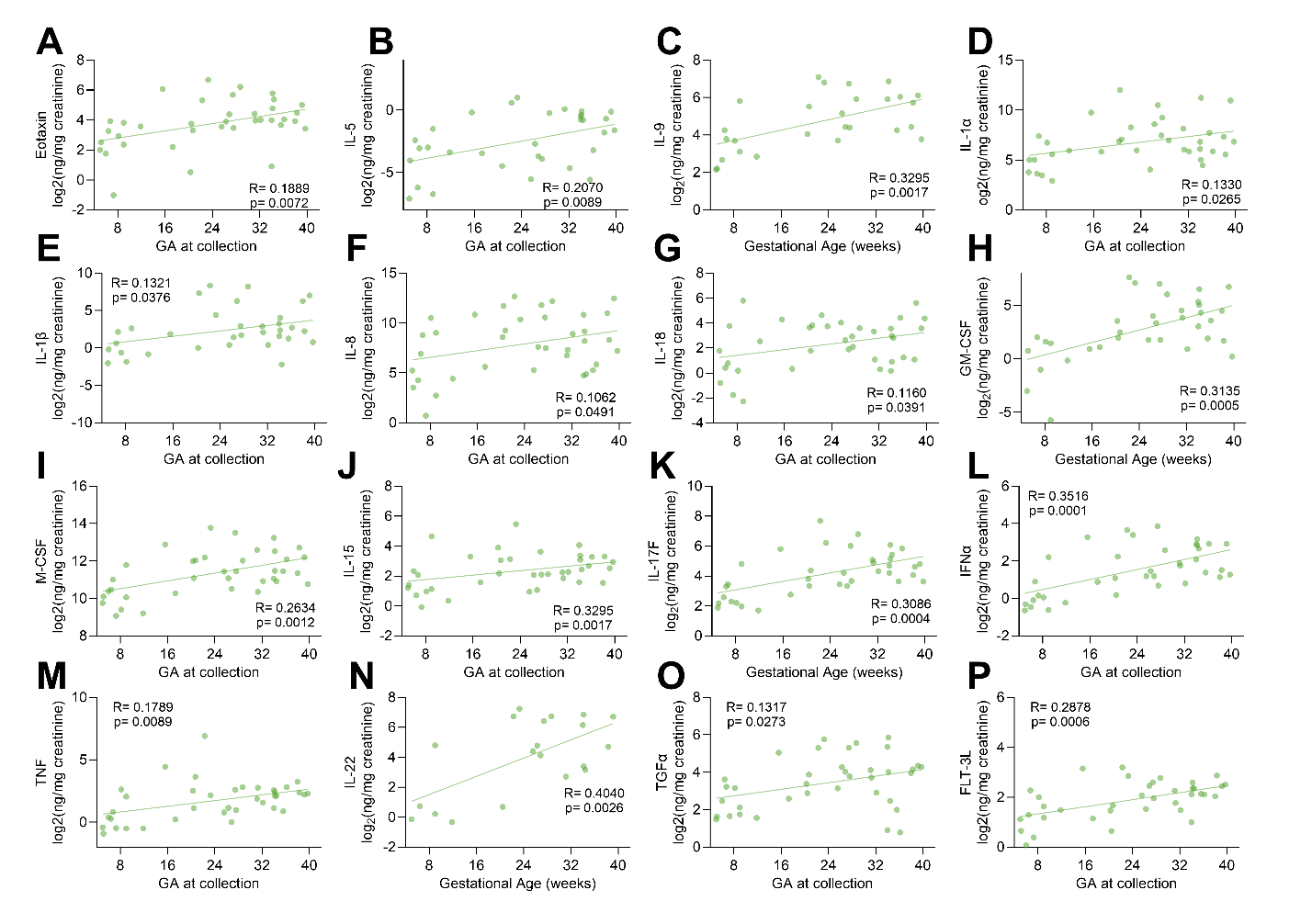

**Supplementary Figure 4. Linear Regression model of urinary cytokines with gestational age in samples with positive bacterial culture.** Linear correlation of eotaxin (**A**), IL-5 (**B**), IL-9 (**C**), IL-1α (**D**), IL-1β (**E**), IL-8 (**F**), IL-18 (**G**), GM-CSF (**H**), M-CSF (**I**), IL-15 (**J**), IL-17F (**K**), IFNα (**L**), TNF (**M**), IL-22 (**N**), TGFα (**O**), and FLT-3L (**P**). *n*=37 (A,D,F,G,I,J,K,L,M,O,P), *n*=32 (B), *n*=27 (C), *n*=33 (E), *n*=35 (H), *n*=20 (N). Data was analyzed by simple linear regression.

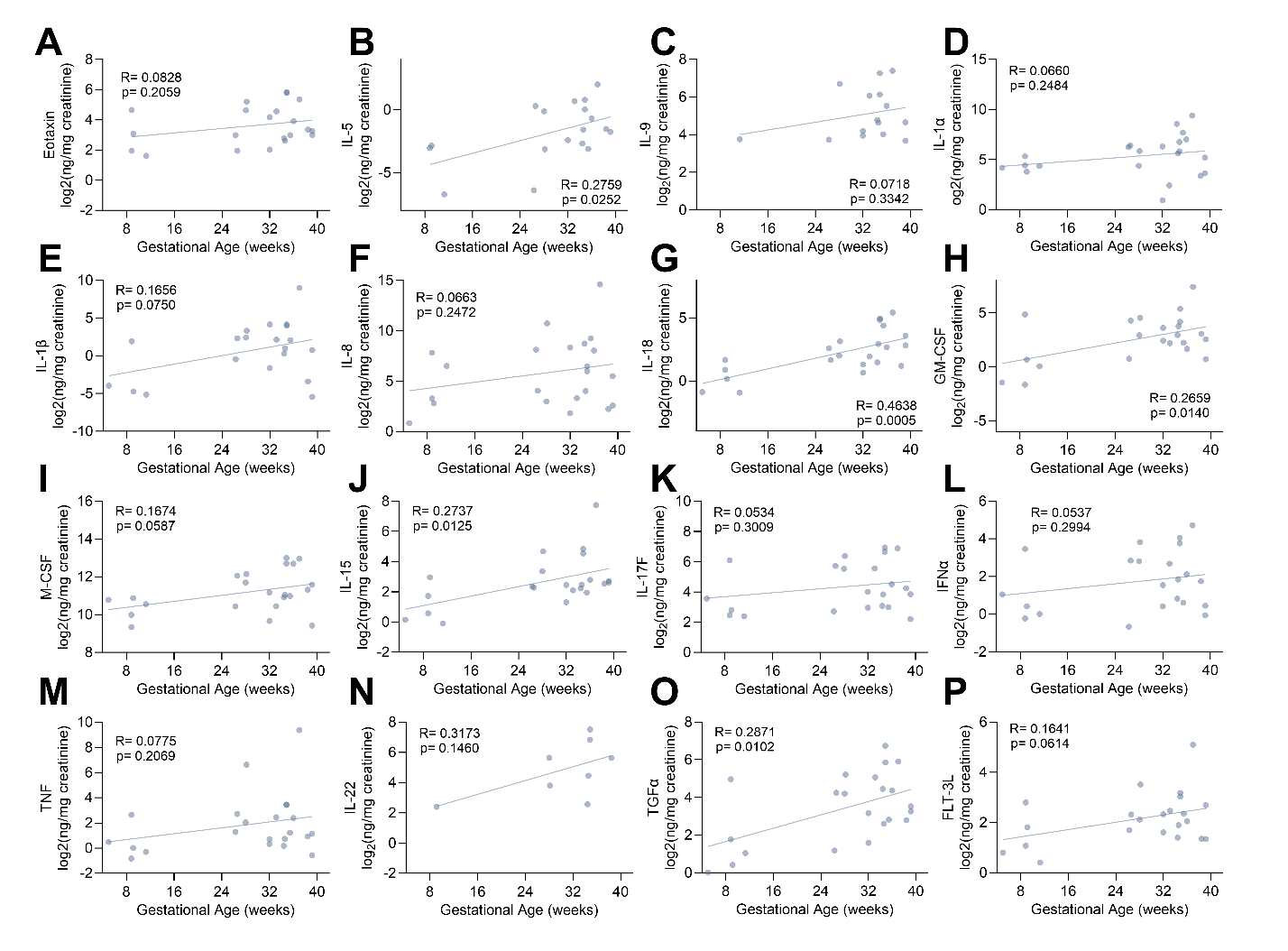

**Supplementary Figure 5. Linear Regression model of urinary cytokines with gestational age in samples with negative bacterial culture.** Linear correlation of eotaxin (**A**), IL-5 (**B**), IL-9 (**C**), IL-1α (**D**), IL-1β (**E**), IL-8 (**F**), IL-18 (**G**), GM-CSF (**H**), M-CSF (**I**), IL-15 (**J**), IL-17F (**K**), IFNα (**L**), TNF (**M**), IL-22 (**N**), TGFα (**O**), and FLT-3L (**P**). *n*=21 (A), *n*=18 (A), *n*=15 (C), *n*=22 (D,F,G,H,I,J,K,L,M,O,P), *n*=20 (E), *n*=8 (N). Data was analyzed by simple linear regression.

**Supplemental Table 2. Patient demographics from nonpregnant and pregnant human urine samples.** Supplemental data to Figure 7.

|  | **Nonpregnant** | | **Pregnant** | | |  |  |
| --- | --- | --- | --- | --- | --- | --- | --- |
|  | **Control (N=8)** | **UTI (N=6)** | **Control (N=22)** | **ASB (N=18)** | **UTI (N=19)** |  | **P-value** |
| **Age** |  |  |  |  |  |  |  |
| Median [Min, Max] | 31.0 [19.0, 39.0] | 27.0 [22.0, 34.0] | 27.0 [19.0, 41.0] | 25.0 [20.0, 39.0] | 23.0 [18.0, 42.0] |  | 0.145 |
| **Race** |  |  |  |  |  |  |  |
| AA | 3 (37.5%) | 2 (33.3%) | 14 (63.6%) | 14 (77.8%) | 8 (42.1%) |  | 0.259 |
| Asian | 1 (12.5%) | 0 (0%) | 1 (4.5%) | 0 (0%) | 0 (0%) |  |  |
| White | 4 (50.0%) | 3 (50.0%) | 5 (22.7%) | 3 (16.7%) | 10 (52.6%) |  |  |
| Other | 0 (0%) | 1 (16.7%) | 2 (9.1%) | 1 (5.6%) | 1 (5.3%) |  |  |
| **Ethnicity** |  |  |  |  |  |  |  |
| Hispanic | 2 (25.0%) | 1 (16.7%) | 3 (13.6%) | 1 (5.6%) | 0 (0%) |  | 0.0321 |
| Not hispanic | 6 (75.0%) | 4 (66.7%) | 19 (86.4%) | 17 (94.4%) | 19 (100%) |  |  |
| Unknown | 0 (0%) | 1 (16.7%) | 0 (0%) | 0 (0%) | 0 (0%) |  |  |
| **BMI** |  |  |  |  |  |  |  |
| Median [Min, Max] | 27.1 [19.3, 34.3] | 23.0 [19.4, 38.4] | 30.5 [23.1, 45.5] | 30.4 [20.7, 44.2] | 29.0 [17.7, 44.9] |  | 0.181 |
| Missing | 0 (0%) | 0 (0%) | 0 (0%) | 1 (5.6%) | 0 (0%) |  |  |
| **Pathogen** |  |  |  |  |  |  |  |
| E. coli | 0 (0%) | 3 (50.0%) | 0 (0%) | 1 (5.6%) | 11 (57.9%) |  | <0.001 |
| S. agalactiae | 0 (0%) | 0 (0%) | 1 (4.5%) | 2 (11.1%) | 3 (15.8%) |  |  |
| Other | 0 (0%) | 3 (50.0%) | 0 (0%) | 0 (0%) | 2 (10.5%) |  |  |
| Mixed perineal flora | 2 (25.0%) | 0 (0%) | 6 (27.3%) | 15 (83.3%) | 1 (5.3%) |  |  |
| Not detected | 6 (75.0%) | 0 (0%) | 15 (68.2%) | 0 (0%) | 2 (10.5%) |  |  |
| **rUTI** |  |  |  |  |  |  |  |
| no | 6 (75.0%) | 4 (66.7%) | 19 (86.4%) | 14 (77.8%) | 11 (57.9%) |  | 0.33 |
| yes | 2 (25.0%) | 2 (33.3%) | 3 (13.6%) | 4 (22.2%) | 8 (42.1%) |  |  |

**Supplemental Table 3. Obstetrical and perinatal information on pregnant human urine samples.** Supplemental data to Figure 8.

|  | **Control (N=22)** | **ASB (N=18)** | **UTI (N=19)** | **P-value** |
| --- | --- | --- | --- | --- |
| **Gestational Age at Collection (weeks)** |  |  |  |  |
| Median [Min, Max] | 32.6 [5.00, 39.1] | 26.6 [5.14, 39.7] | 26.3 [5.00, 38.7] | 0.422 |
| **Gestational Age at Birth (weeks)** |  |  |  |  |
| Median [Min, Max] | 36.7 [29.0, 39.7] | 38.4 [31.7, 39.7] | 37.1 [34.1, 39.9] | 0.238 |
| Missing | 5 (22.7%) | 7 (38.9%) | 7 (36.8%) |  |
| **Preterm** |  |  |  |  |
| yes | 9 (40.9%) | 2 (11.1%) | 5 (26.3%) | 0.332 |
| no | 7 (31.8%) | 9 (50.0%) | 7 (36.8%) |  |
| ND | 6 (27.3%) | 7 (38.9%) | 7 (36.8%) |  |
| **Fetal Sex** |  |  |  |  |
| F | 7 (31.8%) | 3 (16.7%) | 6 (31.6%) | 0.558 |
| M | 10 (45.5%) | 7 (38.9%) | 6 (31.6%) |  |
| ND | 5 (22.7%) | 8 (44.4%) | 7 (36.8%) |  |
| **Fetal Weight at Birth (kg)** |  |  |  |  |
| Median [Min, Max] | 2.90 [1.10, 4.00] | 3.06 [2.27, 3.84] | 2.78 [2.18, 3.29] | 0.541 |
| Missing | 5 (22.7%) | 8 (44.4%) | 9 (47.4%) |  |
